# IL-27 limits HSPC differentiation during infection and protects from stem cell exhaustion

**DOI:** 10.1101/2025.01.15.633135

**Authors:** Daniel L. Aldridge, Zachary Lanzar, Anthony T. Phan, David A. Christian, Ryan Pardy, Booki Min, Ross M. Kedl, Christopher A. Hunter

## Abstract

Many inflammatory stimuli can induce progenitor cells in the bone marrow to produce increased numbers of myeloid cells as part of the process of emergency myelopoiesis. These events are associated with trained immunity and have long-term impacts on hematopoietic stem and progenitor cell (HSPC) development but can also compromise their function. While many cytokines support emergency myelopoiesis, less is known about the mechanisms that temper these events. When mice that lack the cytokine IL-27 were infected with *Toxoplasma gondii*, there was enhanced generation of monocyte progenitors and increased numbers of inflammatory monocytes. In the bone marrow of infected mice there was increased production of IL-27 that localized with HSPCs and a survey of cytokine receptor expression highlighted that HSPCs were uniquely poised to respond to IL-27. Furthermore, the use of *in vitro* differentiation assays and mixed bone marrow chimeras revealed that HSPCs from IL-27 deficient mice are pre-disposed towards the monocyte lineage. Additional studies highlighted that after infection loss of the IL-27R resulted in reduced HSPC fitness that manifested as reduced proliferative responses and a decreased ability to reconstitute the hematopoietic system. Thus, the ability of IL-27 to act on HSPC provides a regulatory brake on differentiation to limit monocyte induction and preserve HSPC stemness.

## Introduction

In response to inflammatory stimuli, cytokine production can directly affect progenitor populations in the bone marrow leading to alterations in hematopoiesis (Collins et al., 2021). For example, in response to microbial challenges, elevated levels of GM-CSF, IFN-γ, and IL-6 modulate hematopoietic stem and progenitor cell (HSPC), which contain hematopoietic stem cells (HSCs) as well as multipotent progenitors (MPP), differentiation and enhance myelopoiesis (Baldridge et al., 2010; Kaufmann et al., 2018; Khan et al., 2020; Link, 2000; MacNamara et al., 2011a; MacNamara et al., 2011b; Maeda et al., 2005; Matatall et al., 2014; Miner et al., 2021; Mirantes et al., 2014; Reynaud et al., 2011). These events can also lead to long-term epigenetic and metabolic changes in HSCs that forms the basis for the process of trained immunity (Ochando et al., 2023). Thus, many of the cytokines produced during different infections are associated with trained immunity (Kaufmann et al., 2018; Khan et al., 2020). For example, after challenge with *Mycobacterium avium* or SARS-COV-2, IFN-γ and IL-6, respectively, can skew HSCs towards myeloid output and increase the functionality of their downstream progeny (Cheong et al., 2023; Matatall et al., 2014).

While trained immunity is associated with long-term changes to hematopoietic stem cells (HSC), inflammation can also reduce HSC functionality, which can manifest as decreased proliferative and differentiation potential. To mitigate these adverse effects, there are cell-intrinsic mechanisms such as autophagic processes to limit oxidative damage, epigenetic reprogramming, and metabolic changes that contribute to the maintenance of stemness (Ibneeva et al., 2024; Liu et al., 2020; Ma et al., 2020; Revuelta and Matheu, 2017; Singh et al., 2018; Zhao and Deininger, 2023). Without these protective mechanisms, HSCs accumulate cellular damage and lose their ability to self-renew and differentiate, a process termed stem cell exhaustion. While inflammation is linked to HSC exhaustion, less is known about the cell extrinsic factors that contribute to the maintenance of HSC quiescence and integrity, but several cytokines have been implicated in this process. For example, in response to repeated stimulation with poly I:C, IFN-α can remove actively proliferating HSCs from the developmental pool as a mechanism to prevent the accumulation of damaged HSCs (Pietras et al., 2014). In addition, during fetal development there is evidence that IL-10 limits HSC responsiveness to inflammatory signaling and thereby limit emergency myelopoiesis as a mechanism to preserve HSC functionality (Collins et al., 2024).

IL-27 is a member of the Type I family of cytokines that utilize JAK-STAT signaling to mediate their biological effects. This family also includes IL-6, and both of these cytokines signal through dimeric receptors that contain the gp130 receptor combined with unique IL-6Rα and IL-27Rα components. Because T cells constitutively express high levels of the IL-27R (Pflanz et al., 2004; Pflanz et al., 2002), the majority of studies on this cytokine have focused on its impact on these populations. Thus, IL-27 can limit the development of pathological Th1, Th2, and Th17 CD4^+^ T cell responses (Findlay et al., 2010; Hamano et al., 2003; Villarino et al., 2003), but several studies have also indicated that the absence IL-27 results in an elevated population of inflammatory monocytes that contribute to the development of immune pathology (Aldridge et al., 2024; Liu et al., 2021). The observation that the loss of IL-27 also results in enhanced inflammatory monocyte induction prior to a detectable T cell response suggests that IL-27 may antagonize the innate events that promote these monocyte populations (Aldridge et al., 2024). Consistent with this idea, there are additional reports that IL-27 can limit neutrophil and monocyte responses and directly impact myelopoiesis (Furusawa et al., 2016; Liu et al., 2014; Liu et al., 2021; Seita et al., 2008; Sun et al., 2017; Wirtz et al., 2006). However, the observation that monocytes do not express the IL-27R suggests that the ability of IL-27 to modulate this population is indirect.

The studies presented here show that infection with *T. gondii* results in increased activity of HPSC and downstream progenitors associated with emergency myelopoiesis; however, in the absence of IL-27, HSPCs display a skewed developmental program that resulted in enhanced monopoiesis. While all HSPCs showed preferential expression of the IL-27R, with LTHSCs and MPP2s exhibiting the highest, the loss of IL-27 during infection resulted in reduced HSPC fitness. Thus, during infection, the ability of IL-27 to act on HSPC provides a regulatory brake on differentiation to limit monocyte induction and preserve HSPC stemness.

## Materials and Methods

### Mice and infection

Female and male C57BL/6 mice were purchased (8-12 weeks old; strain # 000664) from Jackson labs as age and sex matched controls for whole body knockout mice. CD45.1 donor mice were either bred in house or purchased from Jackson labs (10-12 weeks old; strain # 033076). CD45.2 recipient mice were purchased from Taconic labs (10-12 weeks old; model # B6-M). IL-27R^-/-^ (Jackson labs; strain # 018078) were bred in house, as were IL-27p28^-/-^ mice, originally generated by Lexicon Pharmaceuticals, Inc. and previously described (Hall et al., 2012). IL-27-p28-GFP reporters (Kilgore et al., 2018) were similarly bred in house, and the Procr-Ai6 was generated from crossing the Procr-ERT2-cre line (Procr: Jacson labs; strain # 033052) which had been backcrossed 9 generations onto the C57BL/6 background and was provided by Dr. Nancy Speck to the Ai6 mouse line (Jackson labs; strain# 007906). Mice that contain a floxed allele of the IL-27Rα (IL-27R^fl/fl^) were provided by Dr. Booki Min (Do et al., 2017), and maintained within our mouse colony. This line was then crossed with a CD4-cre mouse (Jackson labs; strain# 022071) to generate the CD4^cre^-IL-27R^flox/flox^ mouse line. M1Red mice, which express red fluorescent protein (RFP, encoded by *dsRed2*) under control of the *Irgm1* promoter, were received from Gregory Taylor (Duke University) and maintained in-house. All mice were housed and bred in specific pathogen-free (SPF) facilities in the Department of Pathobiology at the University of Pennsylvania in accordance with institutional guidelines (IACUC# 805045). Cysts of ME49 strain of *T. gondii* were collected from chronically infected CBA mice brain tissues. Experimental mice were then infected i.p. with 20 cysts. For experiments where uninfected bone marrow was sorted, a Prugniaud (PRU) strain of *T. gondii* expressing tdTomato was used (Pru-tdTomato). Both of these are type II strains and show similar virulence.

### Functional Analysis of HSPC *in vivo*

For induction of the Procr-ERT2-cre, tamoxifen was given for three consecutive days (5 mg/mouse) prior to infection. For BRDU proliferation labeling, 5 mg of BRDU was given i.p. to each mouse the same day as infection. 1mg/ml of BRDU was then provided in the drinking water to mice throughout infection. To generate bone marrow chimeras, recipient mice were given a radiation dose of 900-1000 rads at least 6 hrs. prior to transferring donor marrow. Donor marrow was transferred (at least 2×10^6^ cells from each donor, unless otherwise stated; this resulted in a total of 4×10^6^ cells being transferred for mixed chimeras) intravenously (i.v.) via retro-orbital injection. Mice were then given 5 mL of Sulfamethoxazole/ Trimethoprim (200mg+40mg/5ml) in their drinking water and replaced every 2-3 days for two weeks.

### Cell staining, flow cytometry, UMAP analysis, and detection of IL-27

Splenocytes were obtained by grinding spleens through a 70 μm filter and washing with RPMI supplemented with 5% fetal bovine serum (FBS). Red blood cells were then lysed by incubating samples with ACK lysis buffer (Thermo Fischer Scientific) for 5 minutes before washing again with RPMI + 5% FBS. Both femurs and tibias were harvested from mice for bone marrow isolation. Connective tissue was removed from the bones by gentle scraping with a scalpel before one end of each bone was cut off to expose the marrow. Bones were then placed in a 0.5 ml microcentrifuge tube, with a hole punctured in the bottom with a 31g needle. The 0.5ml tube was then placed within a 1.5 ml microcentrifuge tube; the nested tubes were then spun down at 13, 000 rcf for 1.5 min. The pelleted cells were then ACK lysed and washed as above (centrifuging at 500 rcf) before being passed through a 70 μm strainer and resuspended in appropriate media (adapted from (Au - Amend et al., 2016)). For peritoneal exudate cells (PECs), 8 mL of PBS was injected into the peritoneum of mice (21G needle attached to a 10 mL syringe) and re-aspirated using the same needle and syringe. Syringe contents were then expelled into a 15 mL conical tube, spun down at 300 rcf, and resuspended in appropriate media.

After cell numbers were determined for each sample, equivalent numbers of cells were plated in a 96-well plate and spun down at 300 rcf. Cells were then washed with FACS Buffer [1x PBS, 0.2% bovine serum antigen, 1mM EDTA] before incubating with Fc block [99.5 % FACS Buffer, 0.5% normal rat serum, 1 μg/ml 2.4G2 IgG antibody] prior to staining. If CD16/32 was stained for, the staining antibody was substituted for unconjugated 2.4G2 antibody in the Fc block. Cells were stained with the viability dye Ghost Dye Violet 510 (Tonbo biosciences; 12-0870), and the following antibodies were used for subsequent staining: CD90.2 (BD; clone 30-H12), B220 (BD; clone RA3-6B2), Ly6G (BD; clone 1A8), CD11b (BD; clone M1/70), Ly6C (Biolegend; clone HK1.4), Sca-1 (BD/Invitrogen; clone D7), CD117/cKit (Biolegend; clone ACK2), CD135 (Biolegend; clone A2F10), CD150 (Biolegend; clone TC15-12F12.2), CD48 (Biolegend/BD; clone HM48-1), CD127 (Biolegend; clone A7R34), CD16/32 (BD; clone 2.4G2), CD34 (Invitrogen; clone RAM34), CD115 (Biolegend; clone AFS98), CCR2 (Biolegend; clone SA203G11), CX3CR1 (Biolegend; clone SA011F11), CD11c (Biolegend; clone N418), XCR1 (Biolegend; clone ZET), CD172a (Biolegend; clone P84), CD64 (Biolegend; clone X54-5/7.1), F4/80 (Invitrogen; clone BM8), MHCII (Biolegend; clone M5/114.15.2), gp130 (Biolegend; clone 4H1B35), IL-27Rα (BD; 2918), CD132 (BD; clone 4G3), CD131 (BD; clone JORO50), IL-6Rα (Biolegend; clone D7715A7), GM-CSFRα (R&D systems; clone 698423), CD210/IL-10R (Biolegend; clone 1B1.3a), IFN-γR β (Biolegend; clone MOB-47), IFNAR-1 (Biolegend; clone MAR1-5A3), IL-21R (Biolegend; clone 4A9), CD120a/TNFRI (Biolegend; clone 55R-286), CD120b/TNFRII (Biolegend; clone TR75-89), and CD217/IL-17RA (Biolegend; clone S19010F).

Intracellular staining for Ki67 (BD; clone B56), and BCL2 (Biolegend; clone BCL/10C4) was performed using the Foxp3/Transcription Factor Staining Buffer Set (eBioscience) according to the manufacturer’s instructions. BRDU staining was done following a previous published protocol (Matatall, Methods Mol. Biol., 2018). Cells within the vasculature were labeled by injecting 3 μg of florescent anti-CD45 antibody (Biolegend; clone 30-F11)/mouse for 3 min prior to euthanasia. Samples were run on a FACSymphony A3 or A5 (BD) and analyzed using the FlowJo Software analysis program (TreeStar). For UMAP analyses, either 500 or 10,000 live cells (for 5 days of culture or 10, respectively) per sample were sampled using the downsampling plug- in (TreeStar) and subsequently concatenated. The concatenated samples were then used for UMAP analysis with the FlowJo plug-in (TreeStar), using Euclidean distances with the nearest neighbor set to 15, minimum distance of 0.5, and number of components set to 2. All compensated parameters were then used in the analysis. After the UMAP analysis was performed, the X-shift plug-in (TreeStar) was used to further analyze each cluster for its defining features (Samusik et al., 2016).

For IL-27p28 detection, mouse IL-27p28/IL-30 Quantikine ELISA kit (M2728; R&D Systems) was used according to the manufacturer’s protocol.

### Immunofluorescence of bone marrow and image analysis

Procr^tdTomato^IL-27p28^GFP^ mice were generated, infected, and long bones harvested at 5 dpi. The bones were then fixed in 4 % PFA for 2.5-4 hrs, washed with PBS 3x, and then decalcified in 0.5mM EDTA for ≥72 hrs before being paraffin embedded. 5 μm sections were taken and placed onto charged microscope slides. For staining, slides were de-paraffinized, rehydrated, and placed in 1mM EDTA (pH 8.0) overnight in a 55°C water bath to recover epitopes prior to staining. Slides were washed in diH_2_O the next day before proceeding with staining. Staining was adapted from (Im et al., 2019). Slides were blocked with 100μl of blocking buffer (1x PBS+0.3% triton X-100+1:200 rat IgG) for 30 min at room temperature. Slides were then washed 3x, for 5 min each with PBS. Endogenous streptavidin and biotin were then blocked using the ReadyProbes Stretavidin/Biotin Blocking solution (Invitrogen) according to the manufacturer’s directions. After washing, GFP was stained for at 1:40 with anti-GFP AF488 (Biolegend; FM264G) overnight at 4℃. Slides were washed 3x with PBS, as above, and then mounted and coverslipped using 15μl Fluoro-Gel (Electron Microsopy Sciences). Slides were then imaged on a Stellaris Falcon (Leica) microscope.

Images were processed and analyzed using the Fiji software package (Schindelin et al., 2012). Images were converted to binary images, manually thresholded, watersheding applied to separate overlapping objects, and positive cells counted using the “analyze particles” tool.

### Statistics

An unpaired, Welch’s t-test or one-way ANOVA with Sidak’s correction for multiple comparisons were used to test for significant differences between groups. These were used as indicated in each figure and unless otherwise stated. P values of less that 0.05 were considered significant.

## Results

### IL-27 regulates monopoiesis during toxoplasmosis

Infection with *T. gondii* leads to increased production of monocytes that is enhanced in the absence of IL-27 (Aldridge et al., 2024), but it is unclear if this is a result of differentiation from HSPCs or from the more differentiated monocyte progenitors (MPs). Therefore, the Procr-ERT2-cre mouse line, which expresses Cre in HSPCs (Gur-Cohen et al., 2015; Iwasaki et al., 2010), was crossed with the Ai6 reporter line to provide a lineage tracing model. In these mice, tdTomato is expressed in *procr*^+^ cells and, upon tamoxifen (TAM) treatment, HSPCs will be tdTomato^+^ and GFP^+^ but their differentiated progeny will only express zsGreen (Fig. 1A). For these experiments, Procr-Ai6 mice were treated with TAM, infected, and then at 5 dpi the BM and peripheral cells were analyzed. Consistent with other HSPC lineage tracing models (Chapple et al., 2018; Säwen et al., 2018), in naïve bone marrow only a small proportion (∼5%) of long-term hematopoietic stem cells (LTHSCs) were labeled which corresponded to a low proportion of zsGreen^+^ monocytes in the peritoneum and spleen (Fig. 1B-C). After infection, the proportion of zsGreen^+^ LTHSCs in the marrow nearly doubled (Fig. 1B), and the proportion of monocyte dendritic cell progenitors (MDP) also increased (Supp Fig. 1A). This corresponded to an increase in labeled monocytes and neutrophils in the bone marrow of infected mice (Supp. Fig. 1A). In the periphery, there was also an increased proportion of zsGreen^+^ monocytes in the spleen, while in the peritoneum, the initial site of infection and monocyte recruitment, the proportion of labeled monocytes reached ∼100% (Fig. 1C). These progenitors and their progeny were identified based on the expression of defining surface markers as measured by flow cytometry (Supp. Fig. 2). Additional phenotyping revealed that both classical (CCR2^hi^, CX3CR1^lo^) and non-classical (CCR2^lo^, CX3CR1^hi^) monocyte population were labeled to equivalent degrees, indicating that infection impacts the recruitment of both subsets (Supp. Fig 1B). These data sets indicate that the increased myelopoiesis that occurs in response to infection with *T. gondii* is due to expansion and enhanced differentiation of HSPCs, and not just MPs, into downstream progeny, eventually resulting in increased monocyte production.

**Figure 1.**
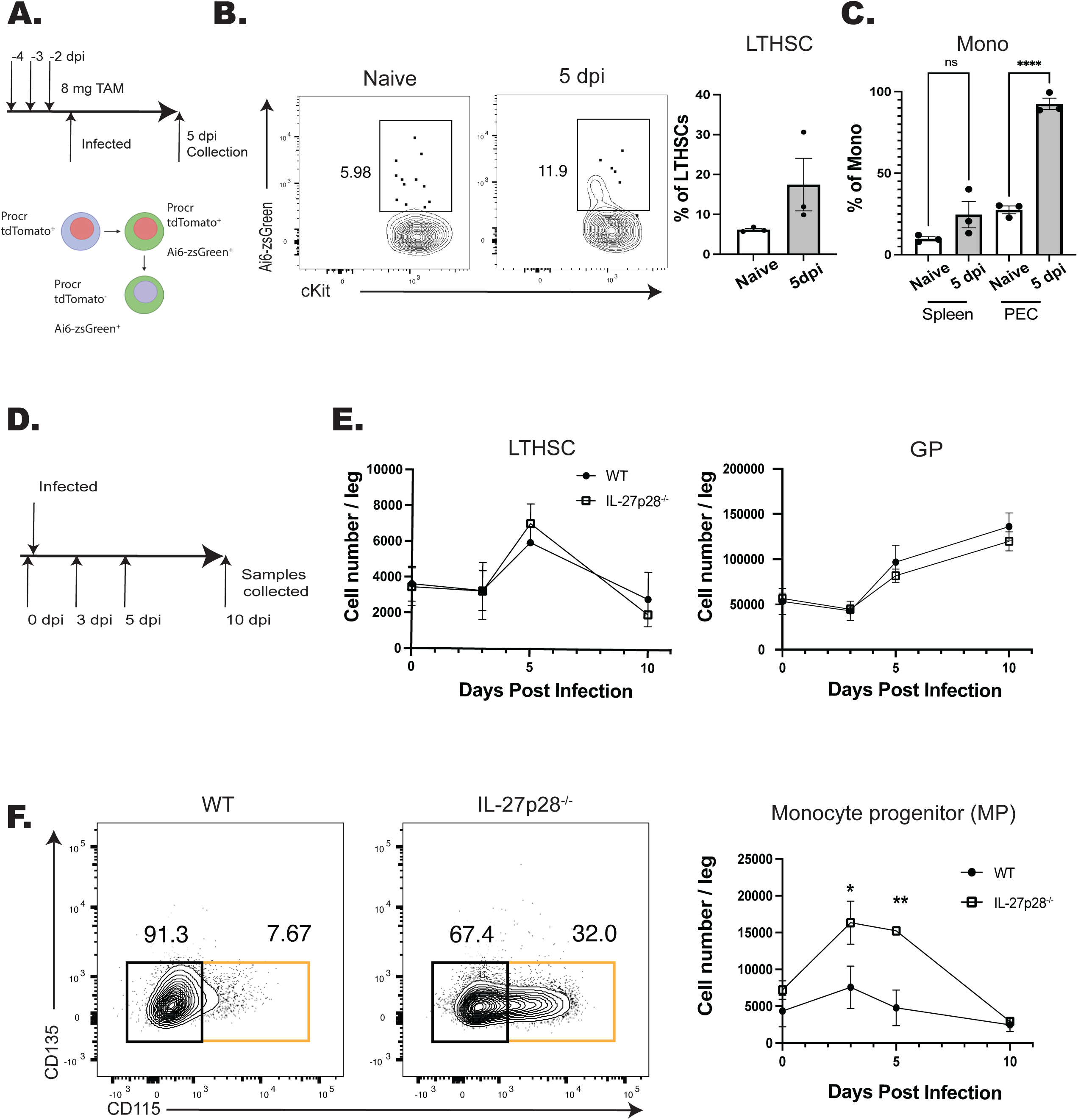
IL-27 regulates monopoiesis during infection. A) Schematic of infection and TAM dosing strategy for Procr-Ai6 mice. B) Procr-Ai6 mice were infected and representative flow plots of LTHSCs (Lin (CD3, NK1.1, B220, Ly6G)^-^, Sca-1^+^, CD117 (cKit)^+^, CD135^-^, CD48^-^, CD150^+^) from naïve (left) and infected (right) mice are shown. Proportions of zsGreen^+^ LTHSCs were then quantified in both groups. C) Proportions of zsGreen^+^ monocytes (CD3^-^, B220^-^, CD11b^+^, Ly6C^+^, Ly6G^-^) in the spleen and PECs of naïve and infected mice. D) Schematic of infection of WT and IL-27p28^-/-^ mice for E-F. E) Numbers of LTHSCs and granulocyte progenitors (GPs) (CD3^-^, NK1.1^-^, B220^-^, CD117^+^, CD34^+^, CD16/32^hi^, Ly6C^+^, CD135^-^, CD115^-^) in the bone marrow of WT and IL-27p28^-/-^ infected mice throughout infection. F) Monocyte progenitors (MPs) (CD3^-^, NK1.1^-^, B220^-^, CD117^+^, CD34^+^, CD16/32^hi^, Ly6C^+^, CD135^-^, CD115^+^; orange box) in the bone marrow of infected WT and IL-27p28^-/-^ mice throughout infection. Representative flow plots are shown (left) and quantified (right). Statistical significance was tested by one-way ANOVA with Sidak’s correction. *, **, and *** correspond to p-values ≤ 0.05, 0.01, and 0.001, respectively. N=3-5 mice/group and data shown are representative of 2-3 experiments.

To determine the impact of IL-27 on this infection induced myelopoiesis, WT and IL-27p28^-/-^ mice were challenged with *T. gondii* and numbers of progenitors (from LTHSCs to MPs) were analyzed (Fig. 1D). While infection led to an expansion of LTHSCs, the numbers of LTHSC, granulocyte progenitors (GP), and other progenitors (including common lymphocyte progenitors (CLPs), common myeloid progenitors (CMPs), and granulocyte monocyte progenitors (GMPs)) (data not shown) were not impacted by the absence of IL-27 (Fig. 1E). Infection also resulted in a transient increase in MPs in WT mice, but in the absence of IL-27 this was enhanced at 3 and 5 dpi before returning to baseline at 10 dpi (Fig. 1F). A similar pattern was observed for the MDPs (Supp. Fig. 1C), and in the IL-27p28^-/-^ mice this corresponded to an increase in mature monocytes in the liver (Supp. Fig. 1D). The CCR2^hi^CX3CR1^lo^ phenotype of these cells indicated that these were inflammatory monocytes, that were predominantly within the tissue parenchyma (Supp. Fig. 1E), suggesting that these monocytes are poised to participate in the tissue pathology associated with these mice. This enhanced monocyte differentiation appeared independent of IFNγ, as blockade of IL-27 during infection in IFNγ reporter mice showed no alteration in cytokine levels in the bone marrow (Supp. Fig. 1F). Thus, one function of IL-27 during acute toxoplasmosis is to limit monopoiesis and the generation of inflammatory monocytes.

### IL-27 regulates monocyte development in a cell intrinsic manner

Previous studies have shown that during toxoplasmosis or trypanosomiasis the absence of IL-27 results in enhanced CD4^+^ T cell responses which contribute to increased accumulation of inflammatory monocytes (Aldridge et al., 2024; Liu et al., 2021). To test if the early (5 dpi) enhanced monopoiesis in the IL-27^-/-^ mice observed here was a secondary consequence of the CD4^+^ T cell response, a CD4^cre^ IL-27R^fl/fl^ (CD4-IL-27R) mouse line was generated. In these mice IL-27R expression was efficiently deleted in T cells, but other IL-27R expressing cells, such as B cells, maintained expression (Supp. Fig. 3A). When infected with *T. gondii* these mice phenocopied whole body IL-27R^-/-^ mice and developed lethal immune pathology > 10 days post-infection (data not shown). When control and CD4-IL-27R mice were infected and analyzed at 5 dpi there were equivalent numbers of monocyte progenitors (Fig. 2A). Thus, the enhanced monopoiesis observed in the whole-body IL-27p28^-/-^ mice is independent of the effect of IL-27 on CD4^+^ T cells.

**Figure 2.**
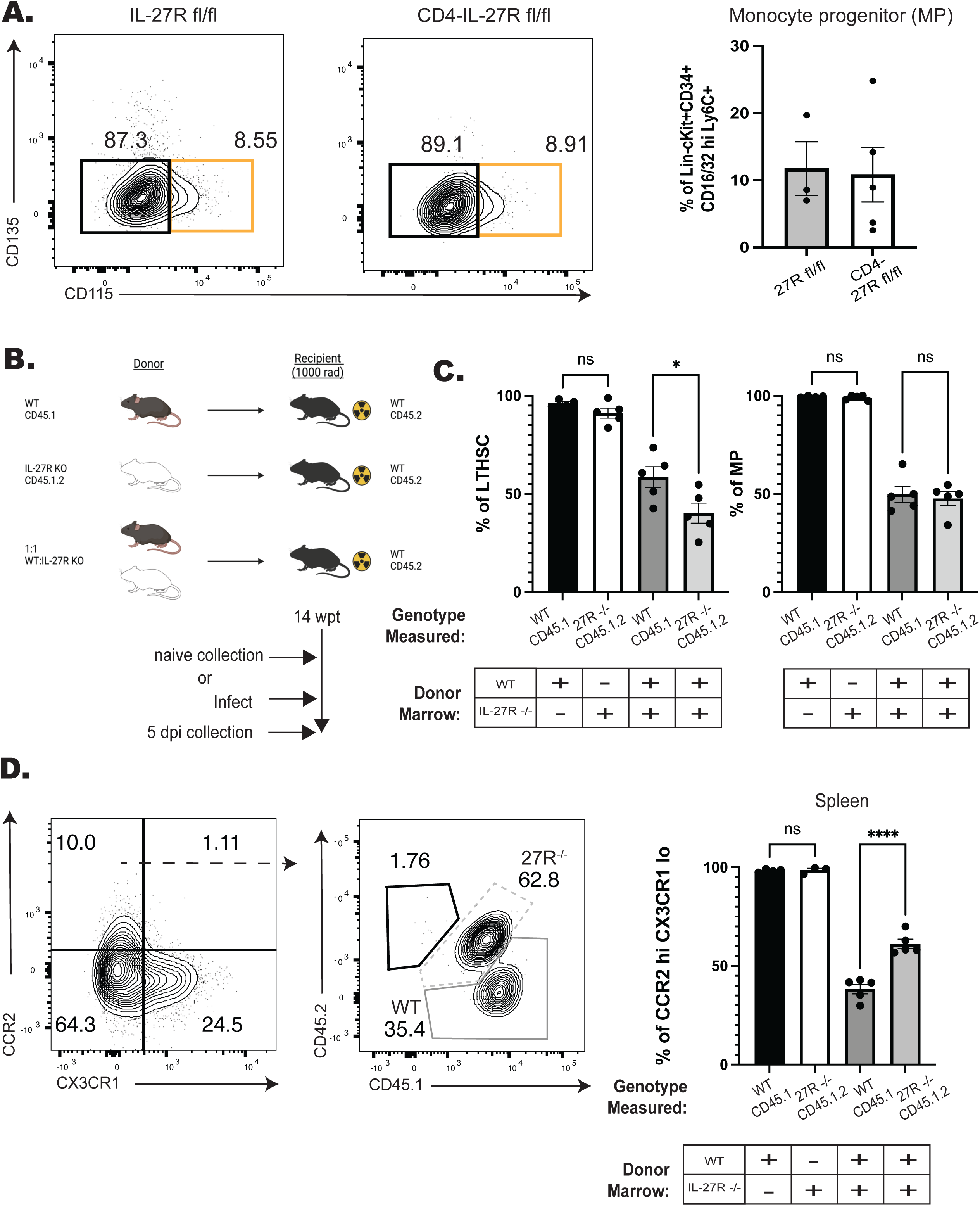
IL-27 regulates monopoiesis during infection in a cell intrinsic manner. A) CD4-IL-27R mice were infected and MPs (orange box) measured at 5 dpi in the BM. Representative flow plots are shown (left) and quantified (right). B) Schematic of BM chimeras generated from WT (CD45.1) and IL-27R^-/-^ (CD45.1.2) bulk bone marrow that were then infected and used in C-D. C) The proportion of LTHSCs (left) and MPs (right) from each respective donor lineage in the bone marrow of both the single (left two bars) and mixed (right two bars) chimeras at 5 dpi. D) Representative flow plots pre-gated on splenic monocytes (left plot) and then down-gated on CCR2^hi^CX3CR1^lo^ (right plot) monocytes at 5 dpi. WT derived monocytes are shown in the solid-dark gray while IL-27R^-/-^ derived cells are in the dashed-light gray. The proportion of each donor lineage that contribute to mature CCR2^hi^CX3CR1^lo^ splenic monocytes at 5 dpi were then quantified (far right). Statistical significance was tested by either Welch’s T-test (A) or one-way ANOVA with Sidak’s correction (C-D). *, **, and *** correspond to p-values ≤ 0.05, 0.01, and 0.001, respectively. N=3-5 mice/group and data shown are representative of 2-3 experiments.

Next, a mixed bone marrow chimera approach was used to determine if the enhanced monopoiesis observed in the absence of IL-27 was cell intrinsic. In these experiments, WT and IL-27R^-/-^ marrow from naïve mice was used to generate a series of individual and mixed bone marrow chimeras (Fig 2B). In the single-transfer chimeras, at fourteen weeks after reconstitution WT and IL-27R^-/-^ marrow reconstituted the hematopoietic system of irradiated recipients with 100% of hematopoietic cells derived from the donor (Supp. Fig. 3B-C, left bars). In the mixed chimeras, while initially no difference in chimerism was observed, as measured by output of peripheral T and B cells, gradually there appeared to be decreased output from the IL-27R^-/-^ lineage. This was confirmed in the bone marrow where a reduced number of IL-27R^-/-^ LTHSCs and downstream MPs was observed (Supp. Fig. 3C, right bars, a 40:60 ratio KO: WT). Despite this reduction in the IL-27R^-/-^ LTHSCs, the population of CCR2^hi^CX3CR1^lo^ monocytes in the periphery were preferentially derived from the IL-27R^-/-^ lineage (Supp. Fig. 3D, right bars, 60:40 ratio KO:WT). To determine if infection would impact this differentiation, these mixed chimeras were infected with *T. gondii* and analyzed at 5 dpi (Fig. 2C). After infection, IL-27R^-/-^ LTHSC remained deficient in the mixed chimeras, but MPs from both lineages were equivalent (Fig. 2C, right bars). Additionally, the CCR2^hi^CX3CR1^lo^ monocytes were still predominately derived from the IL-27R^-/-^ lineage (Fig. 2D, right bars). Together, these data indicate that at homeostasis and during infection IL-27 can support LTHSC populations but also temper the downstream processes that lead to monocyte production. Additionally, this function is independent of IL-27 signals on CD4^+^ T cells, but rather is due to a cell intrinsic ability of IL-27 to limit the production of inflammatory monocytes.

### IL-27p28 is expressed in the bone marrow at homeostasis and during infection

Given the impact of IL-27 on myelopoiesis described above, studies were performed to characterize the cellular sources of IL-27p28 and their spatial location in the bone marrow of naïve and infected mice. WT mice were infected, bone marrow harvested at 0, 3, 5, and 10 dpi, and local production of the IL-27p28 subunit assayed (Fig. 3A). At 0 dpi, low constitutive levels of IL-27p28 were detected and increased throughout infection, with a peak at 5 dpi and elevated levels maintained at 10 dpi (Fig. 3A). The use of an IL-27p28-GFP reporter mouse (Aldridge et al., 2024; Kilgore et al., 2018) revealed a small population of p28-GFP^+^ cells in the bone marrow of naïve mice, but by 5dpi this was expanded (Fig. 3B-C). Additional analysis of the bone marrow and peripheral tissues (spleen, blood, PEC) (Fig. 3B-C) revealed that although a small proportion of T cells were GFP^+^, consistent with other reports (Kimura et al., 2016; Lin et al., 2023), monocytes and macrophages were the dominant source of IL-27 in these sites, by proportion of GFP^+^ cells as well as by cell number (Fig. 3B-C, Supp. Fig. 4A). Additionally, monocyte populations expressed the highest levels of IL-27p28-GFP in comparison to other populations, such as dendritic cells (Supp. Fig. 4B).

**Figure 3.**
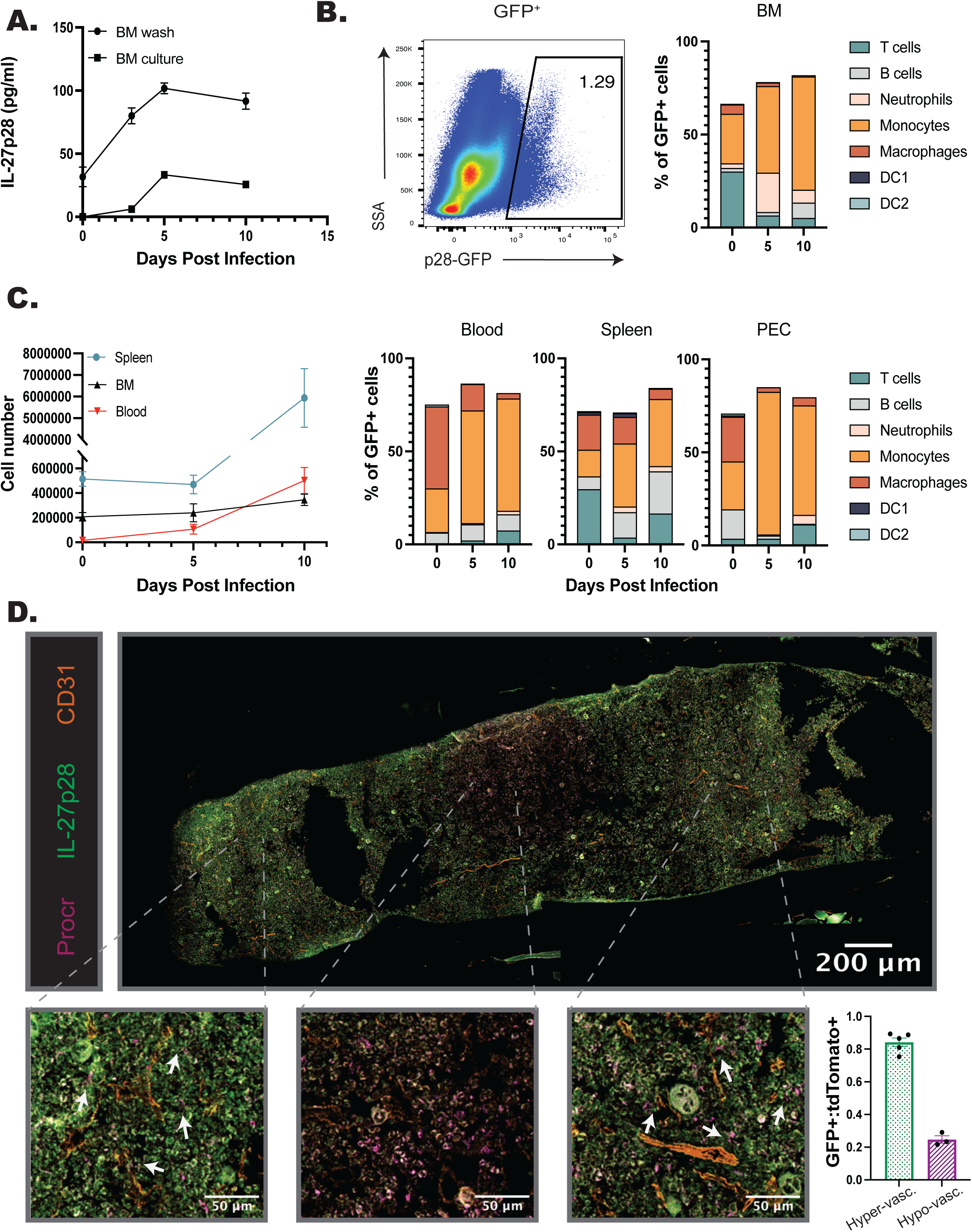
IL-27p28 is produced in the BM during infection. A) WT mice were infected, and the bone marrow harvested throughout infection. BM was either washed with media or cultured for 24 hrs. and supernatant collected. Both the wash and supernatant were analyzed by ELISA for IL-27p28. B) IL-27p28-GFP reporter mice were infected and their bone marrow analyzed by flow cytometry throughout infection. A representative flow plot at 5 dpi of bulk marrow is shown (left) and proportion of immune cell contribution to the GFP^+^ population measured (right). C) Numbers of IL-27p28-GFP^+^ cells in the spleen, blood, and BM of infected mice were quantified throughout infection (left). Proportions of GFP^+^ CD45^+^ cells were analyzed by cell type in the blood, spleen, and PEC during infection (left). D) Femurs from infected Procr-tdTomato^+^p28-GFP^+^ mice were harvested, sectioned, and imaged at 40x magnification. The total imaged femur is shown (top), with zoomed in sections shown below for increased detail of cellular localization (bottom). Arrows indicate areas of localization with Procr-tdTomato^+^ and p28-GFP^+^ cells. Numbers of Procr-tdTomato^+^ and p28-GFP^+^ cells were then counted in eight randomly selected regions throughout the marrow (see Supp. Fig. 4). GFP^+^ cells were normalized to tdTomato^+^ cells and the eight counted regions classified as areas of hyper- or hypo-vascularization according to proximity of the region to the labeled vasculature. This allowed quantification of the localization of cells (bottom right).

To visualize the spatial relationship between IL-27 expressing cells and HSPCs, femurs from naive IL-27p28-GFP mice and imaged. In naïve mice, a stain for Sca-1 revealed low numbers of HSPCs while p28^GFP+^ cells were present at a higher frequency; however, few of the cells in each population colocalized (Supp. Fig. 4C, yellow arrows). During infection the expression of Sca-1 is upregulated by many cell populations (Baldridge et al., 2011; Morales-Mantilla et al., 2022), and so the Procr^tdTomato+^ cre reporter mouse (identifying HSPCs) was crossed with the IL-27p28^GFP^ reporter. As HSCs are typically found within the hypoxic regions of the bone marrow (Eliasson and Jönsson, 2010; Parmar et al., 2007), i.v. injection of anti-CD31 was used to label arterioles to distinguish regions of the marrow that are highly vascularized or more hypoxic. Immunofluorescence of the femur from 5 dpi highlighted that, similar to flow cytometric analysis of the BM of the IL-27p28 reporters, infection resulted in increased numbers of p28^GFP+^ cells (Fig. 3D). Procr^tdtomato^ had a homogenous distribution in the femur (Supp. Fig. 4D, middle) whereas p28^GFP+^ expression was more regional (Fig 3D; Supp. Fig. 4D, top). This was quantified by sampling throughout the femur section (Fig 3D; Supp. Fig. 4D, regions 1-8). The area containing almost exclusively Procr^tdTomato+^ HSPCs, but few p28^GFP+^ cells, was an area of low vascularization, consistent with the HSC niche (Fig. 3D; Supp. Fig. 4D, regions 3-5). However, in regions close to the CD31^+^ vasculature, HSPCs and IL-27p28^+^ cells were found in proximity to one another (Fig. 3D; Supp. Fig. 4D, regions 1-2 and 6-8). When these regions were grouped based on this vascularization, and GFP expressing cells normalized to tdTomato^+^ cells to account for potential random distribution, a marked decrease in GFP^+^ cells was apparent in hypovascularized regions (Fig. 3D). These data indicate that during infection the p28-GFP^+^ BM cells associate with HSPCs, and this is most apparent near the vasculature and away from the HSC niche.

### Developmental expression of the IL-27R by hematopoietic cells

The hematopoietic tree begins with undifferentiated LT-HSCs that differentiate into distinct hematopoietic lineages and mature progeny; the color scheme in Fig. 4A reflects these developmentally related populations. To determine which progenitor stages in the marrow were responsive to IL-27, the expression of the IL-27Rα was compared with other cytokine receptors (gp130, the IL-2R common gamma chain (CD132), IL-6Rα, and GM-CSFRα) that modulate hematopoiesis (Collins et al., 2021; Reynaud et al., 2011). Expression of these receptors was measured on HSPCs and committed progeny in naïve and infected mice (Fig. 4B-C; Fig. 5A-D). The gp130 receptor, one chain of the IL-27 receptor that is also utilized by other IL-6 family members, was expressed by all developmental stages examined in both naïve and infected mice (Fig. 4B). In contrast, the IL-27Rα was highly expressed by LTHSCs, but markedly decreased with lineage commitment, with mature neutrophils and monocytes showing no expression (Fig. 4C). The use of the Haemosphere RNA-seq data set (Choi et al., 2018) and the BloodSpot online tool (Bagger et al., 2015) for the analysis of normal mouse hematopoietic bone marrow transcripts (Chambers et al., 2007; Di Tullio et al., 2011) confirmed that levels of *IL27RA* mRNA were high in LSKs and LTHSCs but decreased with lineage commitment (Supp. Fig. 5A-B). To determine if IL-27R signaling was functional on HSPCs, bulk bone marrow was collected from M1Red reporter mice (Pardy et al., 2024; Stifter et al., 2019), which express RFP under the control of the *Irgm1* promoter, downstream of STAT1 signaling. When these cells were treated with IL-27 (an activator of STAT1) the myeloid committed progenitors showed minimal RFP expression (Supp. Fig. 5C), whereas HSPCS showed a time-dependent increase in reporter expression (Fig. 4D). This differed dramatically from cells isolated *ex vivo* from IRGM1 reporter mice treated with another activator of STAT1, the cytokine IFN-γ. After six hours, all hematopoietic progenitors analyzed exhibited IRGM1-RFP expression (Supp. Fig. 5C and data not shown). The bulk BM cultures stimulated with IL-27 also confirmed that known IL-27R expressing T, B, and NK cells within the culture also responded to IL-27 with a similar kinetic of RFP activation to that of the HSPCs (Supp. Fig. 5C).

**Figure 4.**
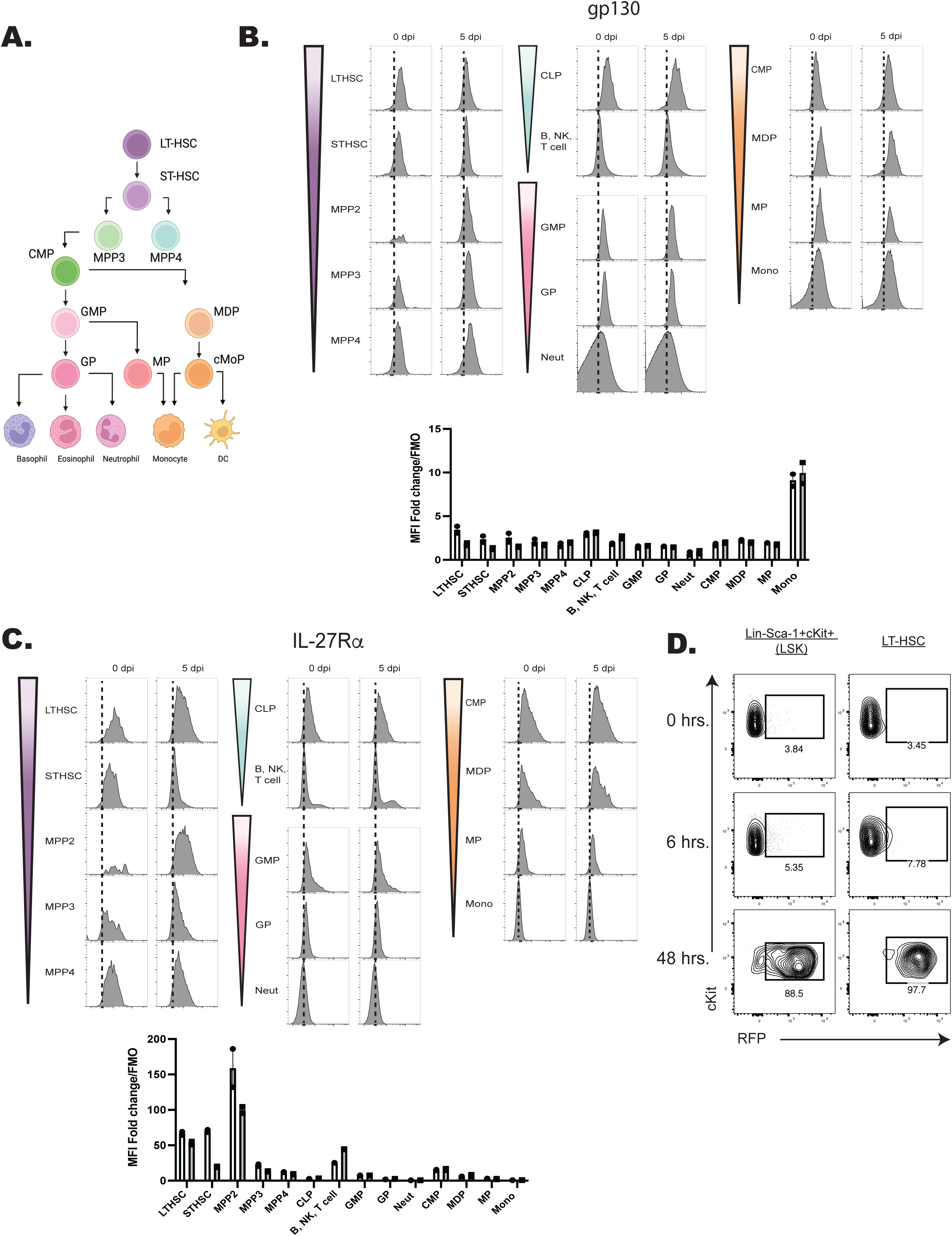
Expression of the IL-27R subunits gp130 and IL-27Rα during hematopoietic development. A) Schematic ball-and-stick model of hierarchical hematopoietic development. Cell type colors are maintained for reference in the following panels. B) Progenitors and progeny shown in (A) were analyzed in WT and infected mice (at 5 dpi) by flow cytometry for the gp130 receptor. 10,000 live cells were concatenated from n=3 mice, and representative flow plots are shown. The fold change of the MFI of the receptor in each cell type over the FMO control for that cell type was then quantified. C) Cells in (B) were analyzed for expression of the IL-27Rα and analyzed as above. D) Bulk bone marrow was isolated from IRGM1-dsRed reporter mice. 22 x 10^6^ cells were plated per condition and stimulated with 20 ng/ml of IL-27 for 0, 6, or 48 hours and RFP expression measured by flow cytometry.

**Figure 5.**
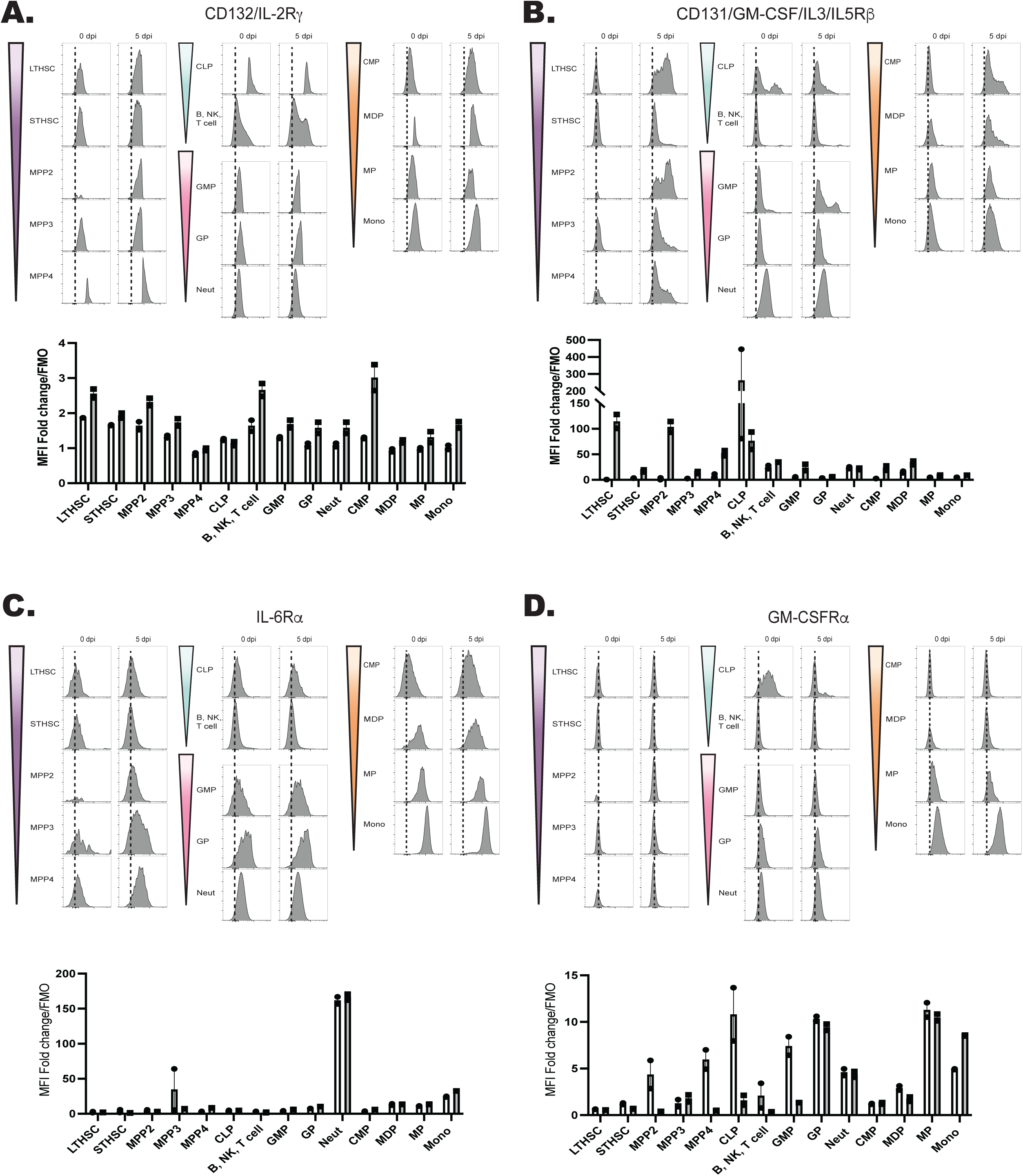
Developmental expression of hematopoietic cytokine receptors. A) Expression of CD132 (IL-2Rγ) and B) CD131 (GM-CSF/IL-3/IL-5Rβ) was analyzed by flow cytometry on progenitors and progeny in the BM of naïve and 5 dpi mice as in Fig. 4. C) Expression of IL-6Rα and D) GM-CSFRα were measured as in (A).

When expression of the common IL-2R gamma chain (utilized by IL-2, −4, −7, −9, −15, and −21), was analyzed, like gp130, it was ubiquitously expressed by all developmental stages, although infection did upregulate its expression (Fig. 5A). Similarly, CD131, which is shared by GM-CSF, IL-3, and IL-5 signaling, was expressed in most developmental stages and was markedly enhanced upon infection (Fig. 5B). In contrast to the IL-27Rα, the expression of IL-6Rα and GM-CSFRα by HSPCs was negligible, and the highest expression of these was detected on fully committed, mature neutrophils and monocytes (Fig. 5C-D). Additionally, both the IL-6Rα and GM-CSFRα showed enhanced expression during infection, further differing from IL-27Rα. A summary of the survey for IL-10R, IFNγRβ, IFNαR, IL-17Rα, TNFIR, and TNFIIR is presented in Supp. Fig. 6, and highlights that while some of these were constitutively expressed across the hematopoietic tree, none of these receptors showed preferential expression by HSPCs. Thus, this survey of cytokine receptor expression highlights that HSPCs but not downstream progenitors are uniquely positioned to respond to IL-27.

### Infection derived IL-27 restrains a monocytic differentiation program in HSPCs

To determine the impact of infection-driven IL-27 on HSPC differentiation, LTHSCs were sorted from naïve or infected WT and IL-27R^-/-^ mice at 5 dpi (see Supp. Fig. 7 for gating strategy). These LTHSCs were then cultured in MethoCult and their phenotype and ability to form colonies was quantified by morphology after 5 or 10 days of culture (Supp. Fig. 8). As expected, cells from naïve mice generated all colony types, but in cultures derived from infected mice there were reduced granulocyte/monocyte (GM) and granulocyte (G) colonies (Supp. Fig. 8A). However, by 10 days post-culture (dpc), there was a slight enhancement in macrophage/monocyte colonies (M) from infected mice. The significant loss of colonies from infected mice, though, made quantification of these colonies difficult. Thus, to better quantify these differences these cultures were analyzed by flow cytometry to measure expression of canonical developmental markers (see materials and methods). Dimensional reduction was then performed via UMAP, and cluster formation was analyzed with X-shift to determine the dominant markers associated with individual clusters. When cells were concatenated from all collection timepoints, 14 clusters were delineated (Supp. Fig. 8B). These clusters where driven, predominantly, by either collection time (Supp. Fig. 8B, top) or proliferative status (Supp. Fig. 8B, bottom). Clusters 1-9 contained only samples from 5 dpc, and clusters 1-6 corresponded to undifferentiated, quiescent colonies that were minimally proliferative (Ki67^lo^) while clusters 7-9 were starting to proliferate (Ki67^hi^) and differentiate to give rise to mature cell populations. By 10 dpc, there was a loss of the quiescent colonies and a rise of populations (clusters 10-14) that are Ki67^hi^ but have increased expression of mature myeloid markers (CD11b, CD16/32, and Ly6C) (Supp. Fig. 8C). This reflects the increased differentiation of colonies into mature myeloid cells. The UMAP analysis highlighted clusters 1, 6, 7, and 14 as the dominant output of this assay (Supp. Fig. 8D): Cluster 1-undifferentiated progenitors (Ki67^lo^, BCL2^mid^, no other expression), Cluster 6-myeloid biased HSPCs (CD150^hi^), Cluster 7-monocyte progenitors (intermediate expression of myeloid markers (CD16/32^lo^), but expression of progenitor markers (CD34^int^)), and cluster 14-mature monocyte (CD11b^hi^, Ly6C^hi^, CD16/32^int^) (Supp. Fig. 8C). Furthermore, this clustering analysis highlighted that infection resulted in a loss of myeloid progenitors (cluster 7) that was further reduced in the absence of IL-27 (Supp. Fig. 8E). However, by 10 dpc this corresponded to enhanced differentiation into mature monocytes (cluster 14).

To compare the ability of HSPCs from WT or IL-27R^-/-^ mice to generate monocyte populations, cells solely from 10 dpc were clustered (Fig. 6A), with the dominant clusters being 1, 2, 3, 10, and 11 (Fig. 6B). Of these, HSPCs were present in cluster 1 (Sca-1^+^cKit/CD117^+^), while cluster 10 contained monocyte progenitors (CD11b^hi^Ly6C^low^CD34^+^CD115^+^), and cluster 11 expressed markers of mature monocytes (CD11b^hi^Ly6C^+^) (Fig. 6C-D). Further analysis (Fig. 6E), revealed that HSPCs from infected mice produced increased numbers of monocytes and macrophages (cluster 11), but only HSPCs from infected, IL-27R^-/-^ mice were enhanced in their differentiation into monocyte progenitors (cluster 10). Thus, during infection the loss of IL-27 signaling enhances the skewing of HSPCs *ex vivo* toward monocyte differentiation.

**Figure 6.**
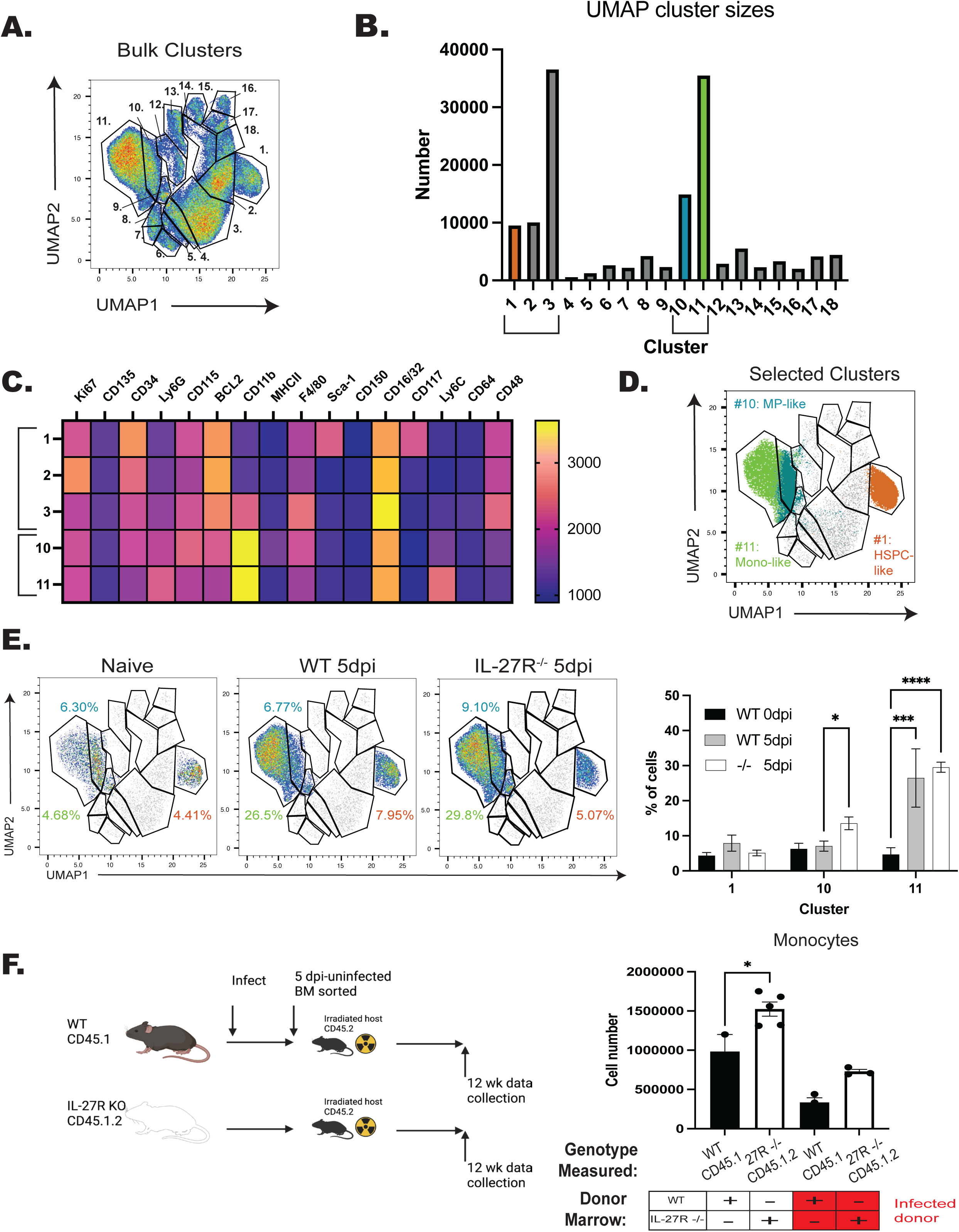
IL-27 limits infection induced HSPC polarization. WT and IL-27R^-/-^ mice were infected for 5 days, LTHSCs sorted, and cultured in MethoCult for 10-12 days before being analyzed by flow cytometry. A) 10,000 live cells from n=3-4 mice/ group (WT (naïve+5 dpi) and IL-27R^-/-^ (5 dpi)) were concatenated and dimensionally reduced via UMAP. B) Numbers of cells in each cluster from (A) were measured. C) X-shift analysis was used to identify the expression level of each marker used in the clustering performed in (A). 5 of the dominant clusters from (B) are shown. D) Clusters 1, 10, and 11, all involved in monocyte development, are highlighted. E) Contribution of Clusters 1, 10, and 11 to each genotype are shown (left) and quantified (right). F) Single bone marrow chimeras were generated from infected WT and IL-27R^-/-^ mice or naïve controls (left). The numbers of single-donor derived, mature monocytes were then measured at 14 weeks post-transplant and compared based on WT vs IL-27R^-/-^ donor (right). Statistical significance was tested by one-way ANOVA with Sidak’s correction. *, ***, and **** correspond to p-values ≤ 0.05, 0.001, and 0.0001, respectively. N=3-5 mice/group and data shown are representative of 2-3 repeated experiments.

In a complimentary *in vivo* approach, single transfers of WT or IL-27R^-/-^ HSPCs from naïve or infected mice were transferred into irradiated WT hosts and differentiation assessed (Fig. 6F). For these experiments bulk marrow was utilized because, as discussed above, inflammation results in upregulation of Sca-1 and other lineage defining markers (Baldridge et al., 2011; Morales-Mantilla et al., 2022), making it difficult to define true HSPCs for transfer by cell-sorting. In addition, for these studies mice were infected with a strain of *T. gondii* that expresses tdTomato (Pru-tdTomato) (Christian et al., 2014; John et al., 2009) to allow parasite removal during sorting and prevent infection of irradiated recipients. Analysis of spleens of recipient mice at 12 weeks, revealed that donors from naive or infected IL-27R^-/-^ mice produced a greater number of monocytes than those derived from WT mice (Fig. 6F). These *ex vivo* and *in vivo* approaches indicate that exposure of LTHSCs to IL-27 during infection results in a stable program that limits monocyte production.

### IL-27 regulates the functionality and fitness of HSPCs during infection

While the data sets presented above indicate a role for IL-27 in the regulation of HSPC differentiation, it was unclear if loss of IL-27 signals during infection would affect their functionality and fitness. To test if there were alteration in HSPC proliferation during infection, WT and IL-27R^-/-^ mice were treated with BRDU throughout infection and HSPC proliferation analyzed at 5 dpi. The combination of Ki67 expression and BRDU incorporation was used to distinguish HSPCs entering cell cycle (Ki67^+^BRDU^-^), cells that have entered cell cycle and are actively proliferating (Ki67^+^BRDU^+^), and those that have proliferated but are no longer in cell cycle (Ki67^-^BRDU^+^). In WT mice, the majority of LSKs are quiescent (Ki67^-^BRDU^-^), a proportion of cells have entered cell cycle (Ki67^+^ BRDU^-^), and a distinct Ki67^+^BRDU^+^ population, as well as a smaller Ki67^-^BRDU^+^ population, can be detected. In the absence of IL-27 signaling, the most prominent difference was a reduced proportion of Ki67^+^BRDU^+^ cells (Fig. 7A). To determine if this decreased Ki67^+^BRDU^+^ population might be due to increased cell death of HSPCs in the IL-27R^-/-^ mice, cells were stained for Annexin V and a free-amine reactive viability dye to discriminate early- and late-apoptotic cells as well as necrotic cells. No difference in cell death was observed between WT and IL-27R^-/-^ HSPCs (Supp. Fig. 9A). These data need to be interpreted with care, as total HSPC numbers are not different between WT and IL-27^-/-^ mice (see Fig 1), but they suggest that IL-27 may regulate either the proliferation or retention of HSPCs within the bone marrow niche.

**Figure 7.**
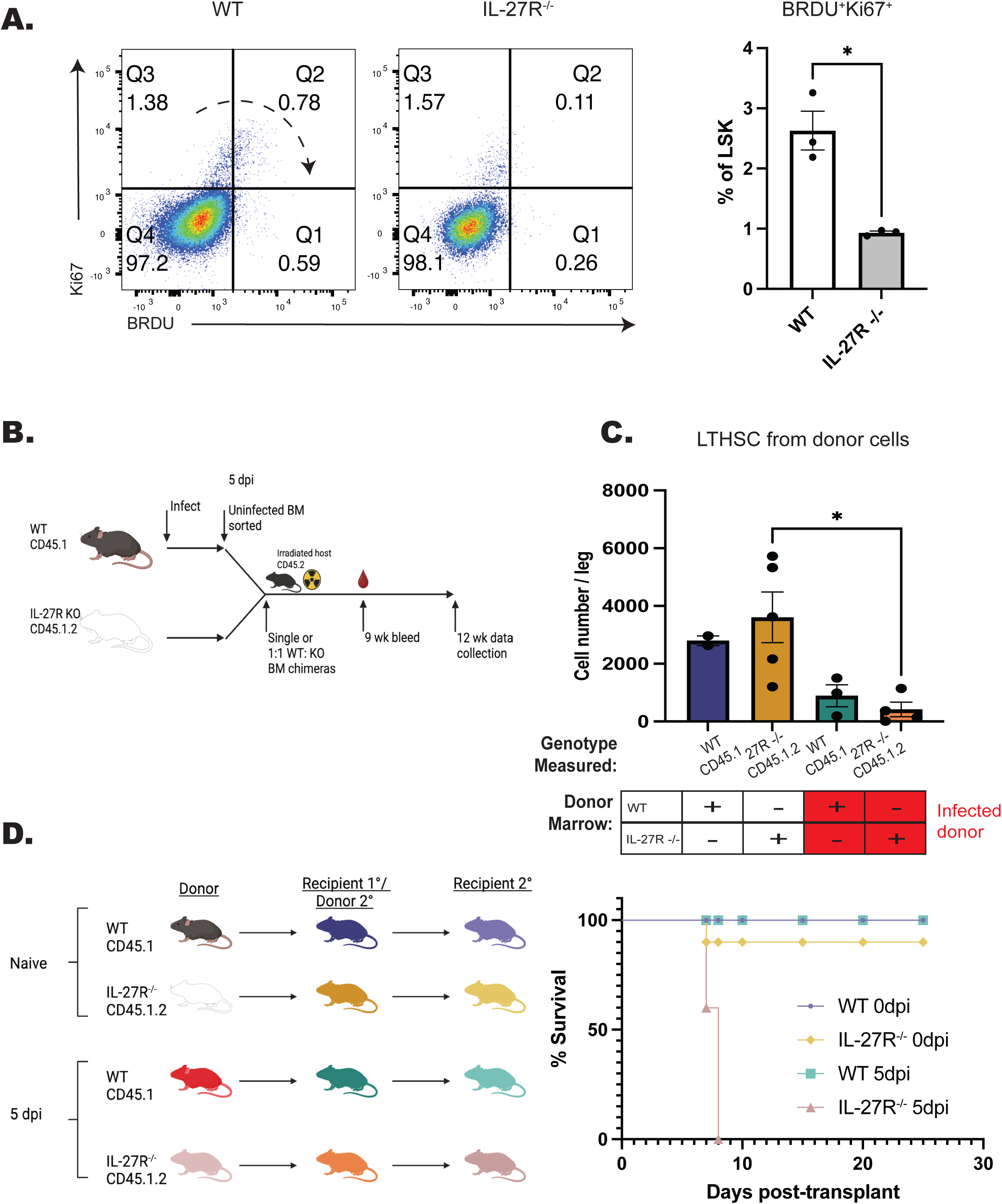
Testing the impact of IL-27 on functionality and fitness of HSPCs during infection. A) WT and IL-27R^-/-^ were infected, treated with 5 mg BRDU, and kept on 1mg/ml BRDU in the drinking water throughout infection. Incorporation of BRDU was then measured in Lin^-^Sca-1^+^cKit^+^ (LSK) progenitor cells in comparison to Ki67 at 5 dpi. Representative flow plots are shown (left) and the proportion of BRDU^+^Ki67^+^ cells quantified (right). B) Schematic for the generation of BM chimeras from infected mice used in (C). C) As in Fig. 6F, numbers of LTHSCs from recipients receiving single-transfer donor marrow from either uninfected or infected mice were measured based on genotype. D) Secondary transplant chimeras from mice that originally received naïve or infected marrow where generated as shown (left). Survival of recipients was then assessed (right).

A key feature of healthy HSCs is their ability to reconstitute the hematopoietic system of irradiated mice, even after serial transfer (Siminovitch et al., 1964; Weissman, 2000). However, inflammation and over-proliferation of HSCs can impair their ability to rescue lethally irradiated recipients (Feng et al., 2008; King et al., 2011; Pietras et al., 2014). To test the impact of IL-27 on HSC fitness and functionality, serial transplantations of HSCs from infected WT and IL-27R^-/-^ mice were performed. In these experiments, marrow from naïve or infected donors was used to produce single chimeras or to provide a 1:1 mix of WT and IL-27R^-/-^ populations that were transferred to irradiated recipients (Fig. 7B). All transfers promoted host survival and at 9 weeks post-transplantation, the WT or IL-27R^-/-^ donors from naïve and infected mice reconstituted the blood compartment with a similar efficiency (Supp. Fig. 9C). However, in the 1:1 chimeras from infected mice, the peripheral compartment was dominated by cells derived from WT mice. Analysis of the marrow of the single transfer chimeras at 12 weeks post-transplant highlighted that when donor cells were derived from infected mice there was a significant reduction in the numbers of LTHSCs (Fig. 7C). To test the functionality of these HSCs, they were then transplanted for a second time into irradiated hosts (Fig. 7D). Mice that received a secondary transplant of marrow from naïve WT and IL-27R^-/-^ mice survived irradiation, consistent with the ability of healthy HSCs to undergo at least three rounds of transplantation (Siminovitch et al., 1964; Weissman, 2000). Additionally, secondary transplantation of WT HSCs from infected mice also resulted in survival of these hosts. However, the serial transfer of marrow from infected IL-27R^-/-^ mice did not save irradiated recipients (Fig. 7D). Thus, while IL-27R^-/-^ HSPCs from infected mice maintain a differentiation bias towards the monocyte lineage, they have a decreased survival and proliferation capacity and a reduced ability to rescue irradiated mice.

## Discussion

During toxoplasmosis, the increased production of inflammatory monocytes is critical to control infection (Dunay et al., 2008), but these populations also have regulatory properties that help to limit immune-pathology. For example, as infection progresses, the presence of IFN-γ in the bone marrow promotes the production of monocytes that produce IL-10 and PGE_2_ (Askenase et al., 2015) which contribute to the restoration of homeostasis. Similarly, during toxoplasmosis and other infections in response to IFN-γ/STAT1 signals (Furusawa et al., 2016 and our own unpublished observations) monocytes become a dominant source of IL-27 (Detavernier et al., 2019; Hall et al., 2012). While IL-27 has a critical role in directly limiting pathological T cell responses, the data presented here show that IL-27 is also part of a regulatory loop that acts on HSPCs to limit emergency myelopoiesis. Infection-induced monopoiesis gives rise to a population of IL-27^+^ monocytes in the bone marrow that can then act on HSPCs to limit further monocyte production. This observation is reminiscent of reports that IFN-γ activated monocytes/macrophages can directly limit HSPCs (McCabe et al., 2015; Seyfried et al., 2020) or the ability of IL-10 to limit emergency myelopoiesis in the fetus (Collins et al., 2024). Likewise, during infection with *T. gondii*, Tregs (Glatman Zaretsky et al., 2017) appear to participate in a regulatory cross-talk with HSPCs. A better understanding of how immune cells in the bone marrow, such as memory B and T, influence HSPCs is needed.

One challenge that faces HSCs during inflammation is the need to protect from stressors associated with proliferation that adversely affect the functions of these cells (Caiado et al., 2021). The concept of HSC “exhaustion” has been developed to describe HSCs that are no longer able to fulfill their role in proliferation and reconstitution of steady-state hematopoiesis (Singh et al., 2020; Zhao and Deininger, 2023). Unanticipated findings from our studies here were the reduced ability of IL-27R^-/-^ HSCs post-infection to compete with WT HSCs, as well as the inability of these HSCs to reconstitute upon secondary transfer, hallmarks of functional exhaustion (Singh et al., 2020). These results infer that the ability of IL-27 to limit emergency myelopoiesis is also associated with the ability of IL-27 to limit HSC exhaustion. It is important to note, however, that these studies indicate that multiple cells within the HSPC category, including MPPs, express the IL-27Rα and are responsive to IL-27. Whether the impacts on HSC exhaustion detected here are due to the direct effects of IL-27 on HSCs or on downstream MPPs, then, warrants further investigation. Aberrant HSPC responses are associated with conditions such as clonal hematopoiesis and aging, and there has been interest in the development of strategies to target these HSPCs. Thus, selective targeting and depletion of “aged” HSPCs via anti-CD150 treatments has been reported to restore balanced hematopoietic output and “youthful” blood phenotypes (Ross et al., 2024). Whether IL-27 treatment can be used to preserve HSPCs, or be blocked to enhance monocyte production, remains to be tested.

A common feature of certain models of trained immunity that affect hematopoiesis is that cytokines, such as IFN-γ and IL-6, promote long term changes that polarize HSPCs toward myeloid differentiation (Cheong et al., 2023; Kaufmann et al., 2018; Khan et al., 2020). Inherent in this process is the idea that HSCs or downstream progeny would be sensitive to different cytokines. The ability to survey the expression of cytokine receptors that influence hematopoiesis emphasized that the shared cytokine receptor chains gp130 and IL-2Rγ are uniformly expressed throughout hematopoietic development. In contrast, the alpha chain receptors for individual cytokines (IL-6Rα, GM-CSFRα, IL-10R, IFNγRβ, IFNAR, IL-21R, TNFRI, and TNFRII) displayed a much more restricted pattern of expression that correlated with key features of the biology of individual cytokines. For example, the limited expression of the IL-6 receptor on downstream progeny is consistent with reports of its high expression on MPPs but not on HSCs (Mirantes et al., 2014; Reynaud et al., 2011). Nevertheless, that HSCs express high levels of the IL-27R, which decreases with differentiation, emphasizes that these stem cells are especially sensitive to the effects of IL-27. Whether IL-27 tempers some of the processes associated with innate training or is itself associated with long-term epigenetic changes is the subject of ongoing studies.

The use of murine models of toxoplasmosis have revealed many of the inhibitory activities of IL-27, and the datasets presented here highlight the ability of IL-27 to influence HSPCs to restrain monocyte/macrophage induction. There are also reports during other infections and models that show IL-27 can regulate neutrophil and monocyte development (Furusawa et al., 2016; Liu et al., 2014; Liu et al., 2021; Seita et al., 2008; Sun et al., 2017; Wirtz et al., 2006). For example, during infection with an attenuated strain of *Plasmodium berghei,* IL-27 was required for the induction of a neutrophil response, and *in vitro* IL-27 blunted the ability of HSPCs to differentiate into the macrophage lineage (Furusawa et al., 2016). However, in less inflammatory settings, such as tumor growth, the overexpression of IL-27 resulted in LSK expansion and the induction of an M1 macrophage population (Zhu et al., 2023). Likewise, in a model of angiotensin-mediated atherosclerosis, IL-27 was needed to overcome HSC quiescence and increase differentiation and output of mature myeloid cells (Peshkova et al., 2019). While it can be difficult to ascribe some of these effects to IL-27 acting directly on HSC, the contrasting nature of these data sets suggest that the impact of IL-27 on HSPCs is context dependent. Indeed, reports that TNFα can upregulate HSPC expression of the IL-27R (He et al., 2020) suggests that IL-27 responsiveness can be regulated by the local environment. A better appreciation of these environmental signals at homeostasis and during different types of inflammation, then, should help to define these context-dependent activities of IL-27 on hematopoiesis.

## Acknowledgments and funding

We would like to thank Dr. Nancy Speck for generously providing us with the Procr-cre mouse line. Additionally, we thank Dr. Gordon Ruthel for his assistance and instruction in capturing images of the mouse bone marrow. Schematic were created using BioRender (agreement license BU27RCNPBH).

Funding for this work was provided by the NIH equipment grant (S10 OD032305-01A1), NIH fellowship support (F31 AI 161962-03), and the Emmerson Foundation.

---

**Supplemental Figure 1.**
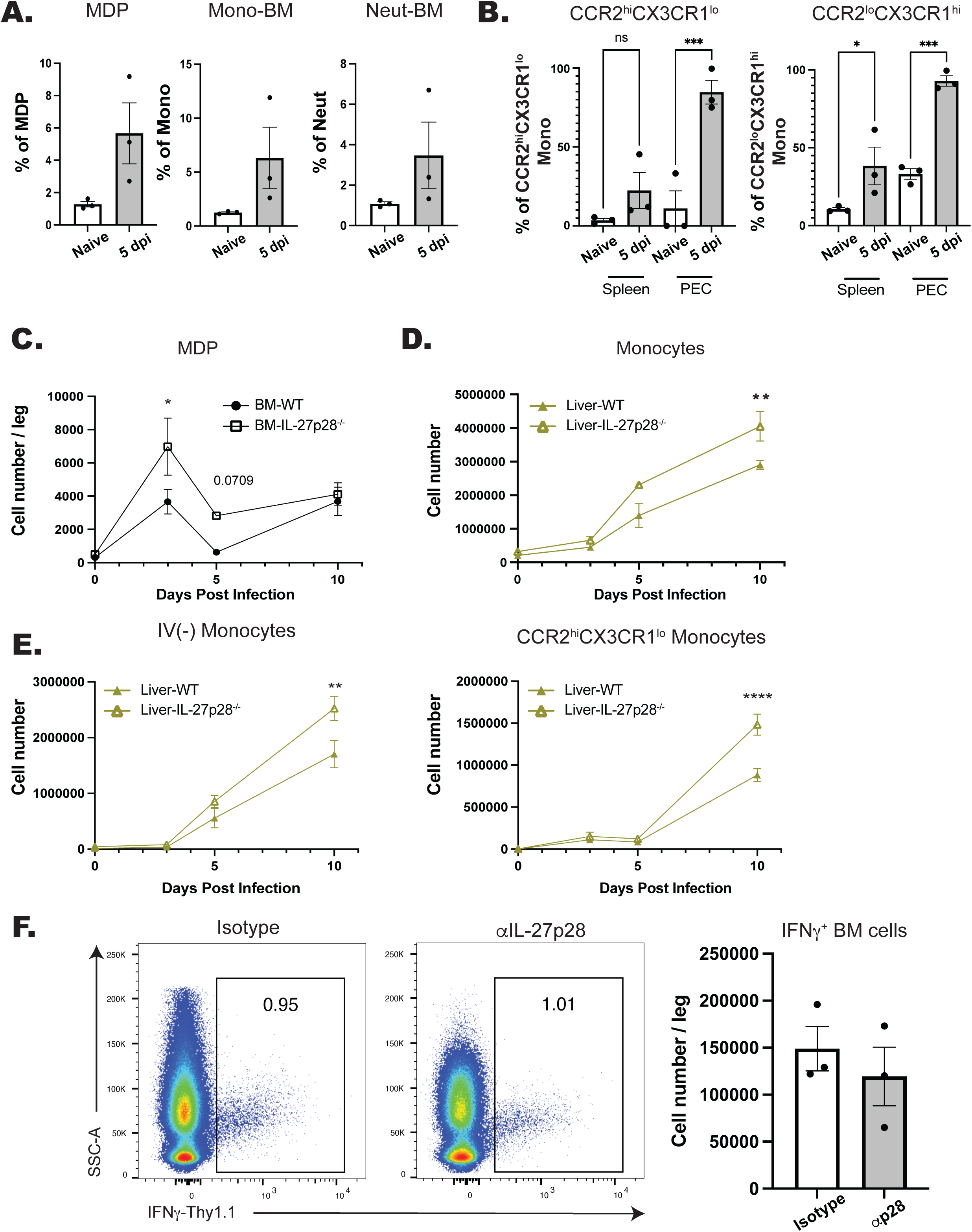
IL-27 regulates monopoiesis during infection. A) The proportion of zsGreen^+^ monocyte dendritic cell progenitors (MDPs) (CD3^-^, NK1.1^-^, B220^-^, CD117^+^, CD34^+^, CD16/32^lo^, CD115^+^), monocytes, and neutrophils (CD3^-^, B220^-^, CD11b^+^, Ly6C^+^, Ly6G^+^) in the BM of naïve and infected Procr-Ai6 mice. B) The proportion of zsGreen^+^ CCR2^hi/lo^ CX3CR1^hi/lo^ monocytes in the spleen and PECs in naïve and infected Procr-Ai6 mice. C) Number of MDPs in WT and IL-27p28^-/-^ mice in the bone marrow during infection. D) Numbers of monocytes in the liver in WT and IL-27p28^-/-^ throughout infection. E) WT and IL-27p28^-/-^ mice were injected I.V. with fluorescent anti-CD45 to label immune cells in the vasculature. Numbers of I.V. label^-^ and CCR2^hi^CX3CR1^lo^ monocytes in the liver were then quantified throughout infection. F) IFNγ-Thy1.1 reporter mice were infected and treated with an anti-p28 blocking antibody. Numbers of IFNγ^+^ cells in the bone marrow was then assessed by flow cytometry. Representative flow plots are shown (left) and quantified (right). Statistical significance was tested by one-way ANOVA with Sidak’s correction. *, **, and *** correspond to p-values ≤ 0.05, 0.01, and 0.001, respectively. N=3-5 mice/group and data shown are representative of 2-3 experiments.

**Supplemental Figure 2.**
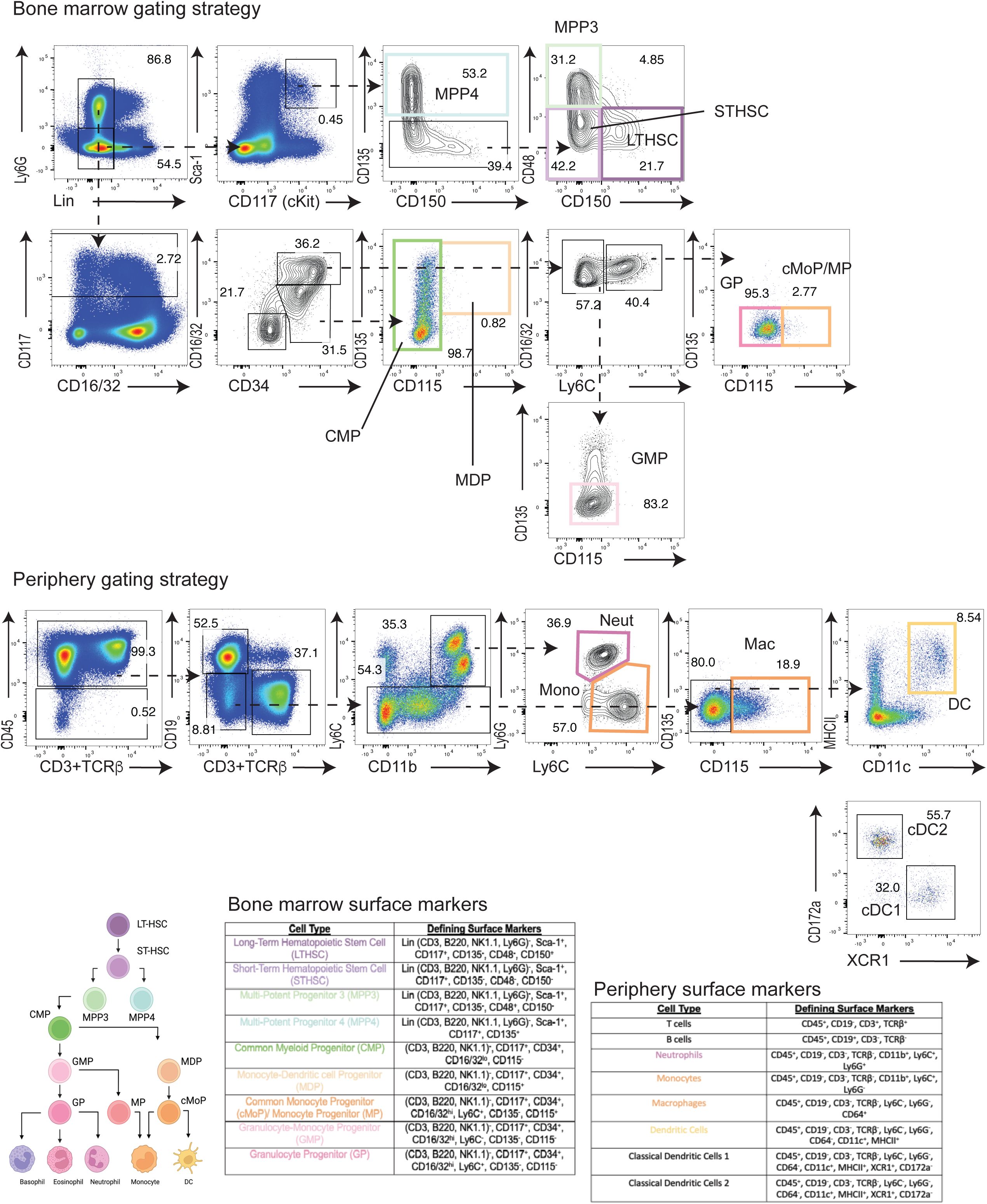
Gating strategy for flow cytometric analysis of cells in the bone marrow and periphery. Representative flow plots from the bone marrow (top) and spleen (bottom) are shown indicating the gating strategy used to identify various cell populations. The combination of surface markers used are listed in the tables (bottom) and are color coded to correspond to both the developmental schematic pictured as well as the corresponding flow gate on the representative plots.

**Supplemental Figure 3.**
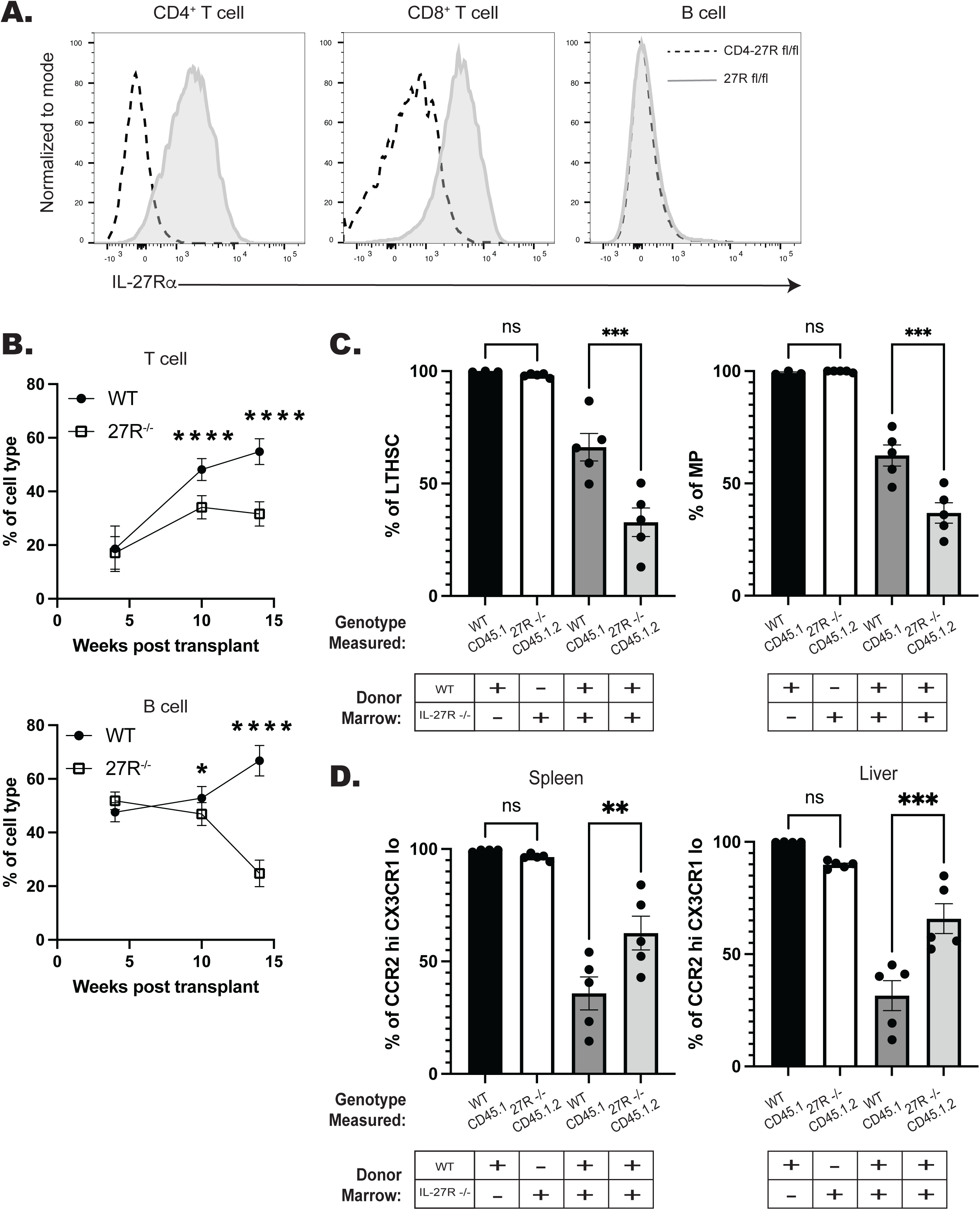
IL-27 regulates monopoiesis during infection and post-irradiation in a cell intrinsic manner. A) IL-27Rα expression on CD4^+^ (CD19^-^CD3^+^CD8α^-^) T cells, CD8^+^ (CD19^-^CD3^+^CD4^-^) T cells, and B cells (CD19^+^CD3^-^) in CD4-27R mice at 5 dpi. B) Reconstitution and chimerism in the peripheral blood 1:1 WT:IL-27R^-/-^ chimeras was measures over time. T (top) and B (bottom) cells from each lineage are shown. C) The proportion of LTHSCs (left) and MPs (right) from each respective donor lineage in the bone marrow post-reconstitution and pre-infection. D) The proportion of CCR2^hi^CX3CR1^lo^ monocytes from each respective donor lineage in the spleen and liver post-reconstitution and pre-infection. Statistical significance was tested by one-way ANOVA with Sidak’s correction. *, **, and *** correspond to p-values ≤ 0.05, 0.01, and 0.001, respectively. N=3-5 mice/group and data shown are representative of 2 repeated experiments.

**Supplemental Figure 4.**
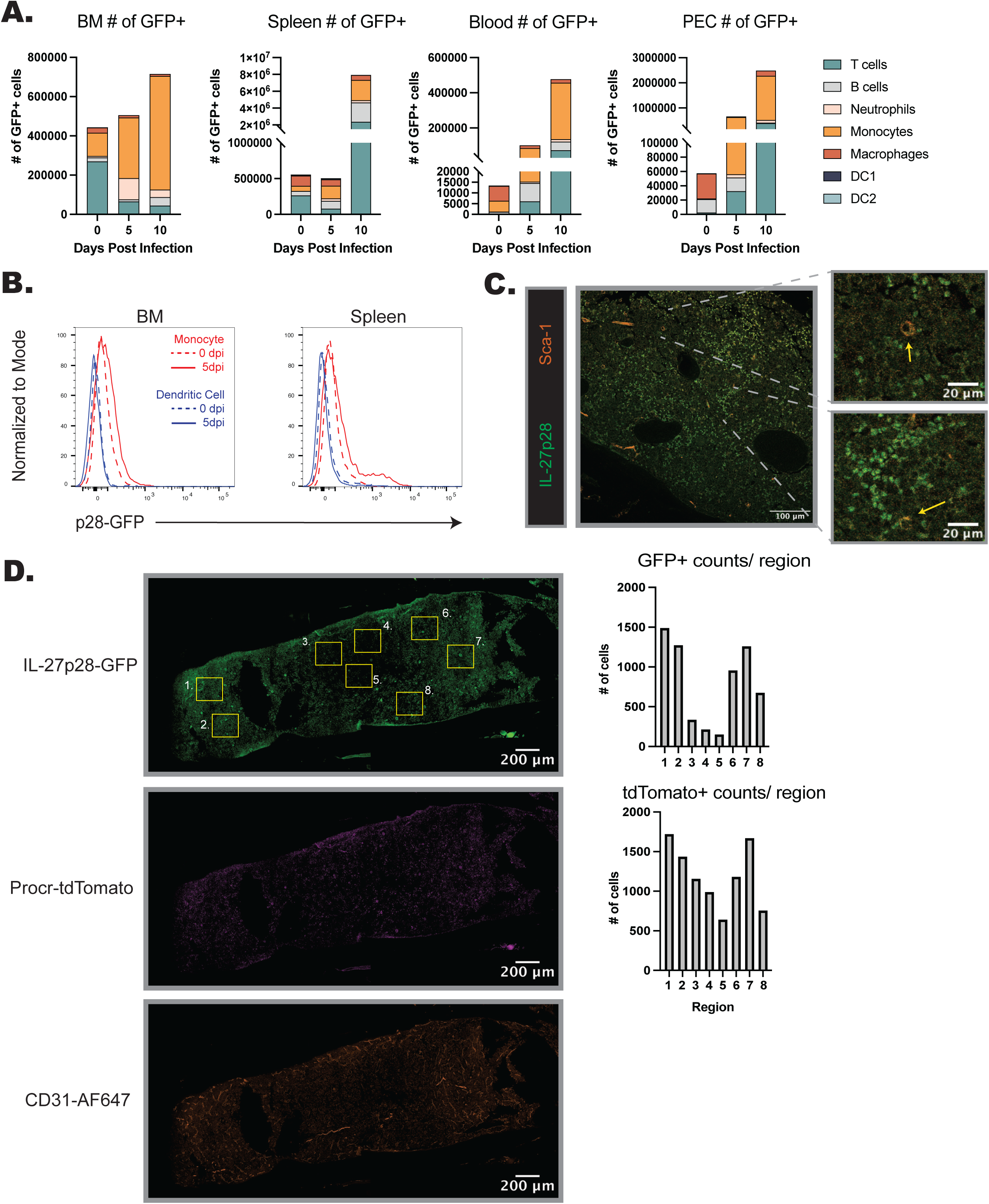
IL-27p28 is produced in the bone marrow during infection. A) IL-27p28-GFP mice were infected, and the numbers of each CD45^+^, immune cell type that expressed GFP in the BM or periphery quantified throughout infection. B) Levels of IL-27p28-GFP expression were measured in bulk monocytes (red) and dendritic cells (blue) in the bone marrow (left) and spleen (right). C) Femurs from naïve IL-27p28-GFP mice were stained for Sca-1 and imaged at 40x magnification. Zoomed in images are shown (right) of the indicated areas, with the yellow arrows indicating positively stained, Sca-1^+^ cells. D) The merged image shown in Fig. 3D is separated by channel (GFP, tdTomato, and AF647, from top to bottom). Eight randomly selected regions that were used for quantification of either GFP^+^ or tdTomato^+^ cells are shown (top), and the quantification of cells in each region is shown next to the corresponding florescence channel (right).

**Supplemental Figure 5.**
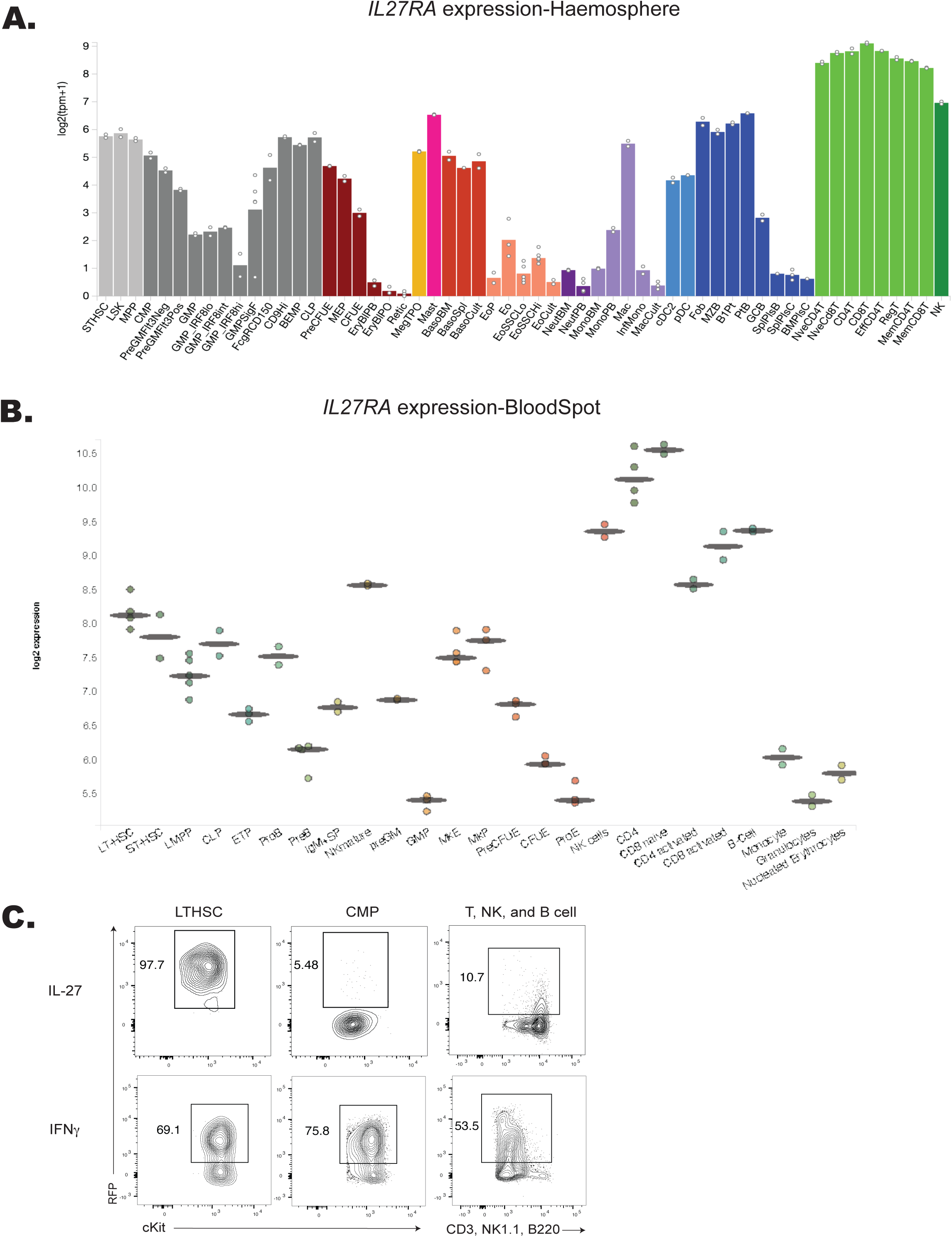
IL-27Rα transcript expression during hematopoiesis and differential sensitivity of LTHSC and CMP to IL-27. A) Expression levels of *IL27RA* are shown from the Haemosphere RNA-seq data set and online tool as well as the Blood Spot data set and visualizer (B). C) Representative flow plots of IRGM1-RFP expression in LTHSCs, CMPs, and lin^+^ T, NK, and B cells 48 hrs. post-stimulation with IL-27 are shown (top). *In vivo* treatment of IRGM1-RFP reporter mice with 1 μg of recombinant IFNγ was performed and all populations analyzed as in (C). Representative flow plots are shown.

**Supplemental Figure 6.**
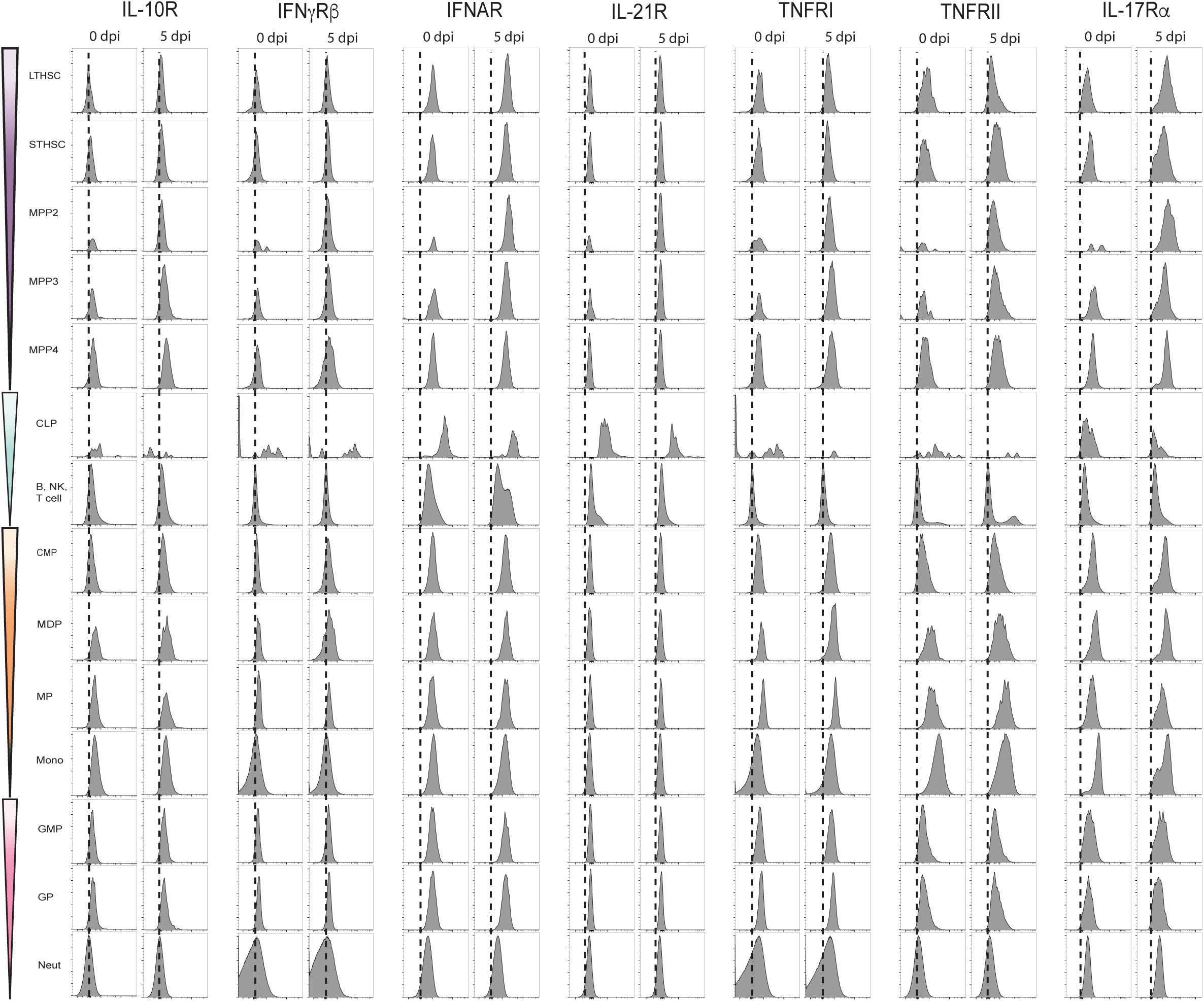
Developmental expression of additional cytokine receptors. Indicated cytokine receptors were analyzed in naive and infected mice as described in Fig. 4-5.

**Supplemental Figure 7.**
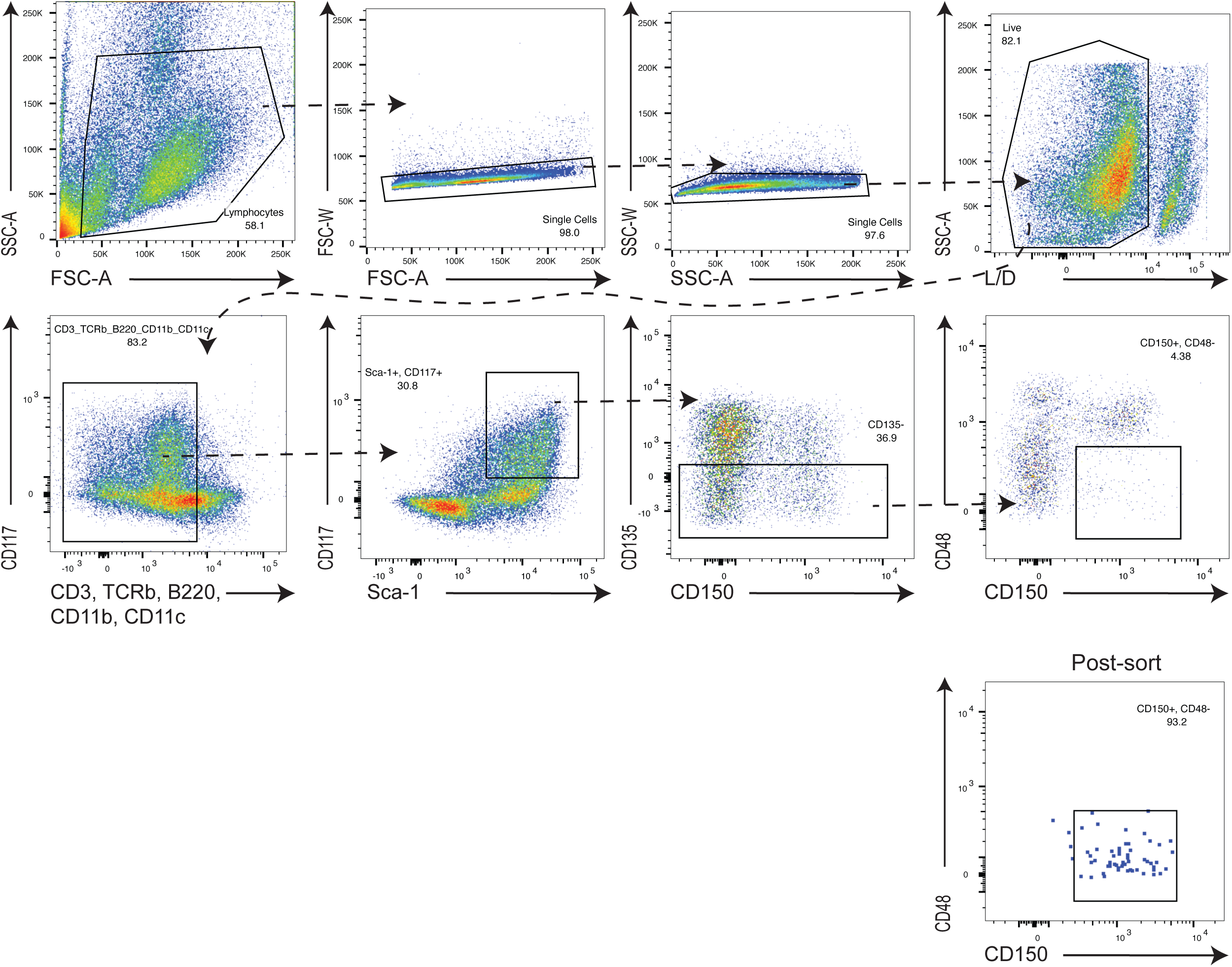
Gating strategy for sorting of LTHSCs used in monocyte differentiation. LTHSCs were sorted from WT naïve, WT 5 dpi, and IL-27R^-/-^ 5 dpi mice. The gating strategy is shown as well as the purity of sorted samples (last panel).

**Supplemental Figure 8.**
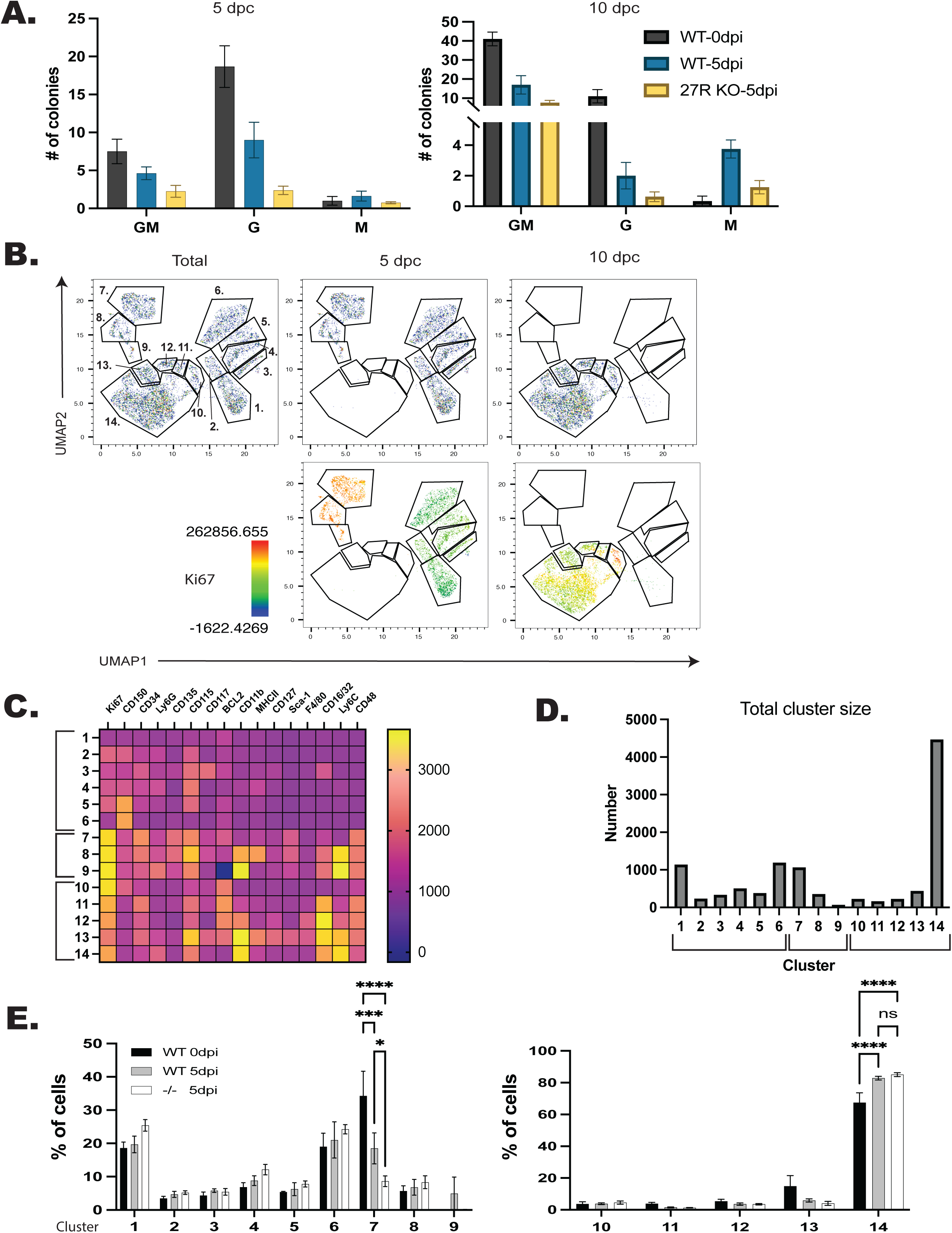
Development and validation of an unbiased flow cytometry approach for the analysis of MethoCult colonies. WT and IL-27R^-/-^ mice were infected, LTHSCs sorted and cultured in MethoCult for either 5 or 10 days. A) Colony phenotypes were analyzed by microscopy and counted by eye. B) Colonies were collected at either 5 or 10 days post-culture (dpc) and analyzed by flow cytometry. 500 live cells from n=3-4 mice/ group were concatenated and dimensionally reduced via UMAP. UMAP plots are shown, and Ki67 expression is overlayed. C) X-shift analysis was used to identify the expression level of each marker used in the clustering performed in (B). D) Numbers of cells in each cluster from (B) were measured. E) Contribution of each genotype to each cluster was quantified. Statistical significance was tested by one-way ANOVA with Sidak’s correction. *, ***, and **** correspond to p-values ≤ 0.05, 0.001, and 0.0001, respectively. N=3-4 mice/group and data shown are from 1 of 3 experiments.

**Supplemental Figure 9.**
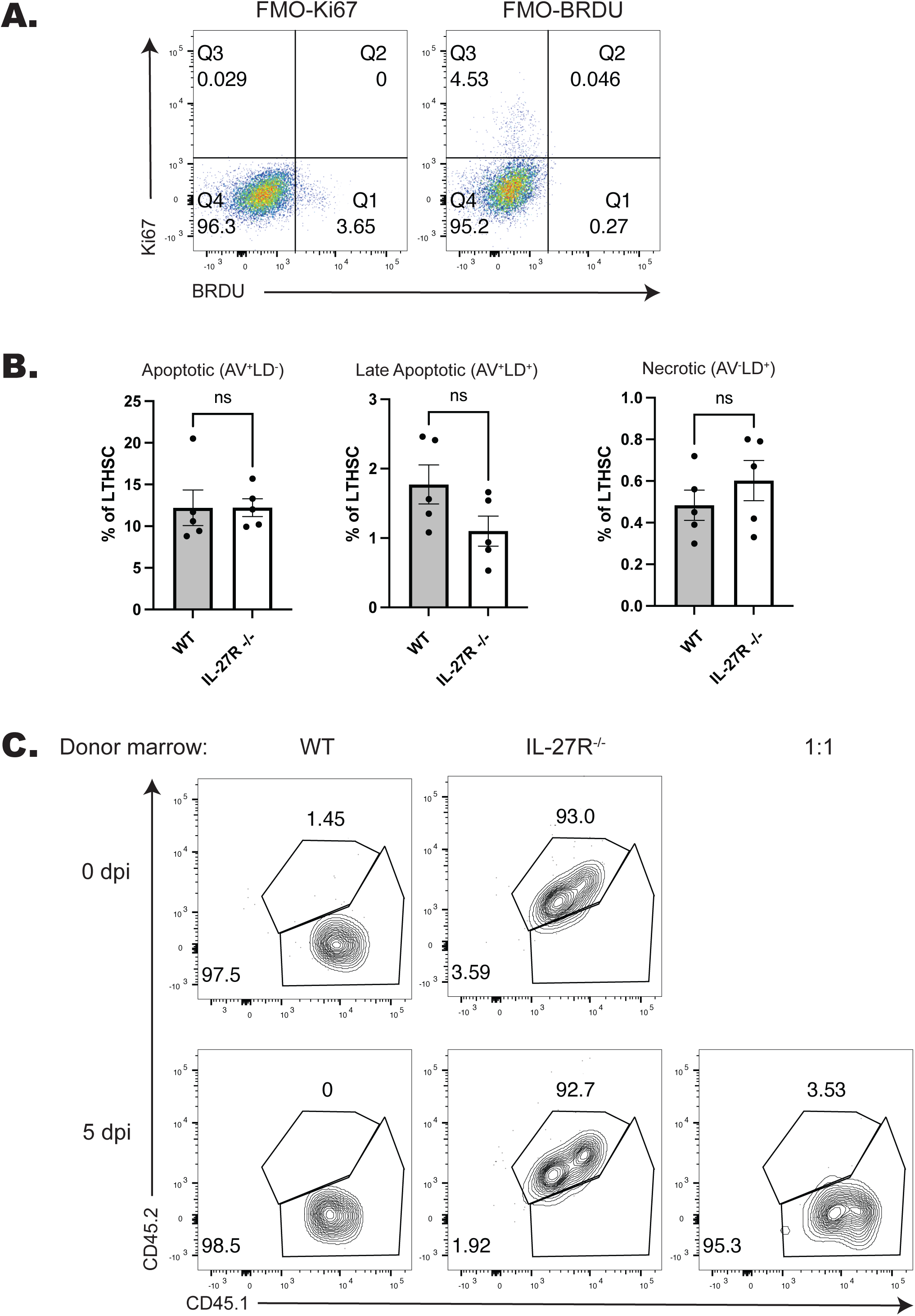
Testing HSPC survival and fitness. A) FMO gating controls for Ki67 and BRDU in the LSK population corresponding to Fig. 7A. B) LTHSCs from WT and IL-27R^-/-^ BM at 5 dpi was stained for Annexin V (AV) expression and free-amine staining viability dye (LD). This allowed the measurement of apoptotic (AV^+^LD^-^), late apoptotic (AV^+^LD^+^), and necrotic (AV^-^LD^+^) cells. The proportion of LTHSCs in each of these stages was quantified. C) Monocyte genotype was analyzed at 9 weeks post-transplant (wpt) from either naïve WT (CD45.1) or IL-27R^-/-^ (CD45.1.2) donors (top) or from infected donors (bottom), including a 1:1 mix of infected WT and IL-27R^-/-^ donors. Statistical significance was tested by Welch’s t-test.

## Notes

### Competing Interest Statement

The authors have declared no competing interest.

### Summary of Updates

In response to reviewer comments at E-life, we have emphasized the expression of the IL-27R on HSPCs and highlight the links to innate trained immunity. New dat sets include expression of IFN-gamma in the bone marrow, improved quantification of cytokine receptor expression during development in the bone marrow and provide additional details of bone marrow reconstitution over time.

## References

Aldridge, D.L., D. Moodley, J. Park, A.T. Phan, M. Rausch, K.F. White, Y. Ren, K. Golin, E. Radaelli, R. Kedl, P.M. Holland, J. Hill, and C.A. Hunter. 2024. Endogenous IL-27 during toxoplasmosis limits early monocyte responses and their inflammatory activation by pathological T cells. mBio 15:e0008324.

Askenase, M.H., S.J. Han, A.L. Byrd, D. Morais da Fonseca, N. Bouladoux, C. Wilhelm, J.E. Konkel, T.W. Hand, N. Lacerda-Queiroz, X.Z. Su, G. Trinchieri, J.R. Grainger, and Y. Belkaid. 2015. Bone-Marrow-Resident NK Cells Prime Monocytes for Regulatory Function during Infection. Immunity 42:1130–1142.

Au - Amend, S.R., K.C. Au - Valkenburg, and K.J. Au - Pienta. 2016. Murine Hind Limb Long Bone Dissection and Bone Marrow Isolation. JoVE e53936.

Bagger, F.O., D. Sasivarevic, S.H. Sohi, L.G. Laursen, S. Pundhir, C.K. Sønderby, O. Winther, N. Rapin, and B.T. Porse. 2015. BloodSpot: a database of gene expression profiles and transcriptional programs for healthy and malignant haematopoiesis. Nucleic Acids Research 44:D917–D924.

Baldridge, M.T., K.Y. King, N.C. Boles, D.C. Weksberg, and M.A. Goodell. 2010. Quiescent haematopoietic stem cells are activated by IFN-gamma in response to chronic infection. Nature 465:793–797.

Baldridge, M.T., K.Y. King, and M.A. Goodell. 2011. Inflammatory signals regulate hematopoietic stem cells. Trends Immunol 32:57–65.

Caiado, F., E.M. Pietras, and M.G. Manz. 2021. Inflammation as a regulator of hematopoietic stem cell function in disease, aging, and clonal selection. Journal of Experimental Medicine 218:

Chambers, S.M., N.C. Boles, K.Y. Lin, M.P. Tierney, T.V. Bowman, S.B. Bradfute, A.J. Chen, A.A. Merchant, O. Sirin, D.C. Weksberg, M.G. Merchant, C.J. Fisk, C.A. Shaw, and M.A. Goodell. 2007. Hematopoietic fingerprints: an expression database of stem cells and their progeny. Cell Stem Cell 1:578–591.

Chapple, R.H., Y.J. Tseng, T. Hu, A. Kitano, M. Takeichi, K.A. Hoegenauer, and D. Nakada. 2018. Lineage tracing of murine adult hematopoietic stem cells reveals active contribution to steady-state hematopoiesis. Blood Adv 2:1220–1228.

Cheong, J.G., A. Ravishankar, S. Sharma, C.N. Parkhurst, S.A. Grassmann, C.K. Wingert, P. Laurent, S. Ma, L. Paddock, I.C. Miranda, E.O. Karakaslar, D. Nehar-Belaid, A. Thibodeau, M.J. Bale, V.K. Kartha, J.K. Yee, M.Y. Mays, C. Jiang, A.W. Daman, A. Martinez de Paz, D. Ahimovic, V. Ramos, A. Lercher, E. Nielsen, S. Alvarez-Mulett, L. Zheng, A. Earl, A. Yallowitz, L. Robbins, E. LaFond, K.L. Weidman, S. Racine-Brzostek, H.S. Yang, D.R. Price, L. Leyre, A.F. Rendeiro, H. Ravichandran, J. Kim, A.C. Borczuk, C.M. Rice, R.B. Jones, E.J. Schenck, R.J. Kaner, A. Chadburn, Z. Zhao, V. Pascual, O. Elemento, R.E. Schwartz, J.D. Buenrostro, R.E. Niec, F.J. Barrat, L. Lief, J.C. Sun, D. Ucar, and S.Z. Josefowicz. 2023. Epigenetic memory of coronavirus infection in innate immune cells and their progenitors. Cell 186:3882–3902.e3824.

Choi, J., T.M. Baldwin, M. Wong, J.E. Bolden, K.A. Fairfax, E.C. Lucas, R. Cole, C. Biben, C. Morgan, K.A. Ramsay, A.P. Ng, M. Kauppi, L.M. Corcoran, W. Shi, N. Wilson, M.J. Wilson, W.S. Alexander, D.J. Hilton, and C.A. de Graaf. 2018. Haemopedia RNA-seq: a database of gene expression during haematopoiesis in mice and humans. Nucleic Acids Research 47:D780–D785.

Christian, D.A., A.A. Koshy, M.A. Reuter, M.R. Betts, J.C. Boothroyd, and C.A. Hunter. 2014. Use of transgenic parasites and host reporters to dissect events that promote interleukin-12 production during toxoplasmosis. Infect Immun 82:4056–4067.

Collins, A., C.A. Mitchell, and E. Passegué. 2021. Inflammatory signaling regulates hematopoietic stem and progenitor cell development and homeostasis. Journal of Experimental Medicine 218:

Collins, A., J.W. Swann, M.A. Proven, C.M. Patel, C.A. Mitchell, M. Kasbekar, P.V. Dellorusso, and E. Passegué. 2024. Maternal inflammation regulates fetal emergency myelopoiesis. Cell 187:1402–1421.e1421.

Detavernier, A., A. Azouz, H. Shehade, M. Splittgerber, L. Van Maele, M. Nguyen, S. Thomas, Y. Achouri, D. Svec, E. Calonne, F. Fuks, G. Oldenhove, and S. Goriely. 2019. Monocytes undergo multi-step differentiation in mice during oral infection by Toxoplasma gondii. Commun Biol 2:472.

Di Tullio, A., T.P. Vu Manh, A. Schubert, G. Castellano, R. Månsson, and T. Graf. 2011. CCAAT/enhancer binding protein alpha (C/EBP(alpha))-induced transdifferentiation of pre-B cells into macrophages involves no overt retrodifferentiation. Proc Natl Acad Sci U S A 108:17016–17021.

Do, J., D. Kim, S. Kim, A. Valentin-Torres, N. Dvorina, E. Jang, V. Nagarajavel, T.M. DeSilva, X. Li, A.H. Ting, D.A.A. Vignali, S.A. Stohlman, W.M. Baldwin, 3rd, and B. Min. 2017. Treg-specific IL-27Rα deletion uncovers a key role for IL-27 in Treg function to control autoimmunity. Proc Natl Acad Sci U S A 114:10190–10195.

Dunay, I.R., R.A. Damatta, B. Fux, R. Presti, S. Greco, M. Colonna, and L.D. Sibley. 2008. Gr1(+) inflammatory monocytes are required for mucosal resistance to the pathogen Toxoplasma gondii. Immunity 29:306–317.

Eliasson, P., and J.I. Jönsson. 2010. The hematopoietic stem cell niche: low in oxygen but a nice place to be. J Cell Physiol 222:17–22.

Feng, C.G., D.C. Weksberg, G.A. Taylor, A. Sher, and M.A. Goodell. 2008. The p47 GTPase Lrg-47 (Irgm1) Links Host Defense and Hematopoietic Stem Cell Proliferation. Cell Stem Cell 2:83–89.

Findlay, E.G., R. Greig, J.S. Stumhofer, J.C. Hafalla, J.B. de Souza, C.J. Saris, C.A. Hunter, E.M. Riley, and K.N. Couper. 2010. Essential role for IL-27 receptor signaling in prevention of Th1-mediated immunopathology during malaria infection. J Immunol 185:2482–2492.

Furusawa, J., I. Mizoguchi, Y. Chiba, M. Hisada, F. Kobayashi, H. Yoshida, S. Nakae, A. Tsuchida, T. Matsumoto, H. Ema, J. Mizuguchi, and T. Yoshimoto. 2016. Promotion of Expansion and Differentiation of Hematopoietic Stem Cells by Interleukin-27 into Myeloid Progenitors to Control Infection in Emergency Myelopoiesis. PLoS Pathog 12:e1005507.

Glatman Zaretsky, A., C. Konradt, F. Dépis, J.B. Wing, R. Goenka, D.G. Atria, J.S. Silver, S. Cho, A.I. Wolf, W.J. Quinn, J.B. Engiles, D.C. Brown, D. Beiting, J. Erikson, D. Allman, M.P. Cancro, S. Sakaguchi, L.F. Lu, C.O. Benoist, and C.A. Hunter. 2017. T Regulatory Cells Support Plasma Cell Populations in the Bone Marrow. Cell Rep 18:1906–1916.

Gur-Cohen, S., T. Itkin, S. Chakrabarty, C. Graf, O. Kollet, A. Ludin, K. Golan, A. Kalinkovich, G. Ledergor, E. Wong, E. Niemeyer, Z. Porat, A. Erez, I. Sagi, C.T. Esmon, W. Ruf, and T. Lapi 2015. PAR1 signaling regulates the retention and recruitment of EPCR-expressing bone marrow hematopoietic stem cells. Nat Med 21:1307–1317.

Hall, A.O., D.P. Beiting, C. Tato, B. John, G. Oldenhove, C.G. Lombana, G.H. Pritchard, J.S. Silver, N. Bouladoux, J.S. Stumhofer, T.H. Harris, J. Grainger, E.D. Wojno, S. Wagage, D.S. Roos, P. Scott, L.A. Turka, S. Cherry, S.L. Reiner, D. Cua, Y. Belkaid, M.M. Elloso, and C.A. Hunter. 2012. The cytokines interleukin 27 and interferon-γ promote distinct Treg cell populations required to limit infection-induced pathology. Immunity 37:511–523.

Hamano, S., K. Himeno, Y. Miyazaki, K. Ishii, A. Yamanaka, A. Takeda, M. Zhang, H. Hisaeda, T.W. Mak, A. Yoshimura, and H. Yoshida. 2003. WSX-1 Is Required for Resistance to Trypanosoma cruzi Infection by Regulation of Proinflammatory Cytokine Production. Immunity 19:657–667.

He, H., P. Xu, X. Zhang, M. Liao, Q. Dong, T. Cong, B. Tang, X. Yang, M. Ye, Y. Chang, W. Liu, X. Wang, Z. Ju, and J. Wang. 2020. Aging-induced IL27Ra signaling impairs hematopoietic stem cells. Blood 136:183–198.

Ibneeva, L., S.P. Singh, A. Sinha, S.E. Eski, R. Wehner, L. Rupp, I. Kovtun, J.A. Pérez-Valencia, A. Gerbaulet, S. Reinhardt, M. Wobus, M. von Bonin, J. Sancho, F. Lund, A. Dahl, M. Schmitz, M. Bornhäuser, T. Chavakis, B. Wielockx, and T. Grinenko. 2024. CD38 promotes hematopoietic stem cell dormancy. PLoS Biol 22:e3002517.

Im, K., S. Mareninov, M.F.P. Diaz, and W.H. Yong. 2019. An Introduction to Performing Immunofluorescence Staining. Methods Mol Biol 1897:299–311.

Iwasaki, H., F. Arai, Y. Kubota, M. Dahl, and T. Suda. 2010. Endothelial protein C receptor-expressing hematopoietic stem cells reside in the perisinusoidal niche in fetal liver. Blood 116:544–553.

John, B., T.H. Harris, E.D. Tait, E.H. Wilson, B. Gregg, L.G. Ng, P. Mrass, D.S. Roos, F. Dzierszinski, W. Weninger, and C.A. Hunter. 2009. Dynamic Imaging of CD8(+) T cells and dendritic cells during infection with Toxoplasma gondii. PLoS Pathog 5:e1000505.

Kaufmann, E., J. Sanz, J.L. Dunn, N. Khan, L.E. Mendonça, A. Pacis, F. Tzelepis, E. Pernet, A. Dumaine, J.C. Grenier, F. Mailhot-Léonard, E. Ahmed, J. Belle, R. Besla, B. Mazer, I.L. King, A. Nijnik, C.S. Robbins, L.B. Barreiro, and M. Divangahi. 2018. BCG Educates Hematopoietic Stem Cells to Generate Protective Innate Immunity against Tuberculosis. Cell 172:176–190.e119.

Khan, N., J. Downey, J. Sanz, E. Kaufmann, B. Blankenhaus, A. Pacis, E. Pernet, E. Ahmed, S. Cardoso, A. Nijnik, B. Mazer, C. Sassetti, M.A. Behr, M.P. Soares, L.B. Barreiro, and M. Divangahi. 2020. M. tuberculosis Reprograms Hematopoietic Stem Cells to Limit Myelopoiesis and Impair Trained Immunity. Cell 183:752–770.e722.

Kilgore, A.M., S. Welsh, E.E. Cheney, A. Chitrakar, T.J. Blain, B.J. Kedl, C.A. Hunter, N.D. Pennock, and R.M. Kedl. 2018. IL-27p28 Production by XCR1(+) Dendritic Cells and Monocytes Effectively Predicts Adjuvant-Elicited CD8(+) T Cell Responses. Immunohorizons 2:1–11.

Kimura, D., M. Miyakoda, K. Kimura, K. Honma, H. Hara, H. Yoshida, and K. Yui. 2016. Interleukin-27-Producing CD4(+) T Cells Regulate Protective Immunity during Malaria Parasite Infection. Immunity 44:672–682.

King, K.Y., M.T. Baldridge, D.C. Weksberg, S.M. Chambers, G.L. Lukov, S. Wu, N.C. Boles, S.Y. Jung, J. Qin, D. Liu, Z. Songyang, N.T. Eissa, G.A. Taylor, and M.A. Goodell. 2011. Irgm1 protects hematopoietic stem cells by negative regulation of IFN signaling. Blood 118:1525–1533.

Lin, C.-H., C.-J. Wu, S. Cho, R. Patkar, W.J. Huth, L.-L. Lin, M.-C. Chen, E. Israelsson, J. Betts, M. Niedzielska, S.A. Patel, H.G. Duong, R.R. Gerner, C.-Y. Hsu, M. Catley, R.A. Maciewicz, H. Chu, M. Raffatellu, J.T. Chang, and L.-F. Lu. 2023. Selective IL-27 production by intestinal regulatory T cells permits gut-specific regulation of TH17 cell immunity. Nature Immunology 24:2108–2120.

Link, D.C. 2000. Mechanisms of granulocyte colony-stimulating factor-induced hematopoietic progenitor-cell mobilization. Semin Hematol 37:25–32.

Liu, F.D.M., E.E. Kenngott, M.F. Schröter, A. Kühl, S. Jennrich, R. Watzlawick, U. Hoffmann, T. Wolff, S. Norley, A. Scheffold, J.S. Stumhofer, C.J.M. Saris, J.M. Schwab, C.A. Hunter, G.F. Debes, and A. Hamann. 2014. Timed Action of IL-27 Protects from Immunopathology while Preserving Defense in Influenza. PLOS Pathogens 10:e1004110.

Liu, G., O. Abas, Y. Fu, Y. Chen, A.B. Strickland, D. Sun, and M. Shi. 2021. IL-27 Negatively Regulates Tip-DC Development during Infection. mBio 12:

Liu, Y., Y. Chen, X. Deng, and J. Zhou. 2020. ATF3 Prevents Stress-Induced Hematopoietic Stem Cell Exhaustion. Front Cell Dev Biol 8:585771.

Ma, Z., J. Xu, L. Wu, J. Wang, Q. Lin, F.A. Chowdhury, M.H.H. Mazumder, G. Hu, X. Li, and W. Du. 2020. Hes1 deficiency causes hematopoietic stem cell exhaustion. Stem Cells 38:756–768.

MacNamara, K.C., M. Jones, O. Martin, and G.M. Winslow. 2011a. Transient activation of hematopoietic stem and progenitor cells by IFNγ during acute bacterial infection. PLoS One 6:e28669.

MacNamara, K.C., K. Oduro, O. Martin, D.D. Jones, M. McLaughlin, K. Choi, D.L. Borjesson, and G.M. Winslow. 2011b. Infection-induced myelopoiesis during intracellular bacterial infection is critically dependent upon IFN-γ signaling. J Immunol 186:1032–1043.

Maeda, K., Y. Baba, Y. Nagai, K. Miyazaki, A. Malykhin, K. Nakamura, P.W. Kincade, N. Sakaguchi, and K.M. Coggeshall. 2005. IL-6 blocks a discrete early step in lymphopoiesis. Blood 106:879–885.

Matatall, K.A., C.C. Shen, G.A. Challen, and K.Y. King. 2014. Type II interferon promotes differentiation of myeloid-biased hematopoietic stem cells. Stem Cells 32:3023–3030.

McCabe, A., Y. Zhang, V. Thai, M. Jones, M.B. Jordan, and K.C. MacNamara. 2015. Macrophage-Lineage Cells Negatively Regulate the Hematopoietic Stem Cell Pool in Response to Interferon Gamma at Steady State and During Infection. Stem Cells 33:2294–2305.

Miner, X., Z. Shanshan, D. Fang, Z. Qingyun, W. Jinhong, W. Chenchen, Z. Caiying, Z. Sen, L. Bingqing, W. Peng, and E. Hideo. 2021. Granulocyte colony-stimulating factor directly acts on mouse lymphoid-biased but not myeloid-biased hematopoietic stem cells. Haematologica 106:1647–1658.

Mirantes, C., E. Passegué, and E.M. Pietras. 2014. Pro-inflammatory cytokines: emerging players regulating HSC function in normal and diseased hematopoiesis. Exp Cell Res 329:248–254.

Morales-Mantilla, D.E., B. Kain, D. Le, A.R. Flores, S. Paust, and K.Y. King. 2022. Hematopoietic stem and progenitor cells improve survival from sepsis by boosting immunomodulatory cells. eLife 11:e74561.

Ochando, J., W.J.M. Mulder, J.C. Madsen, M.G. Netea, and R. Duivenvoorden. 2023. Trained immunity — basic concepts and contributions to immunopathology. Nature Reviews Nephrology 19:23–37.

Pardy, R.D., K.A. Walzer, B.A. Wallbank, J.H. Byerly, K.M. O’Dea, I.S. Cohn, B.E. Haskins, J.L. Roncaioli, E.J. Smith, G.Y. Buenconsejo, B. Striepen, and C.A. Hunter. 2024. Analysis of intestinal epithelial cell responses to Cryptosporidium highlights the temporal effects of IFN-γ on parasite restriction. PLOS Pathogens 20:e1011820.

Parmar, K., P. Mauch, J.A. Vergilio, R. Sackstein, and J.D. Down. 2007. Distribution of hematopoietic stem cells in the bone marrow according to regional hypoxia. Proc Natl Acad Sci U S A 104:5431–5436.

Peshkova, I.O., T. Aghayev, A.R. Fatkhullina, P. Makhov, E.K. Titerina, S. Eguchi, Y.F. Tan, A.V. Kossenkov, M.V. Khoreva, L.V. Gankovskaya, S.M. Sykes, and E.K. Koltsova. 2019. IL-27 receptor-regulated stress myelopoiesis drives abdominal aortic aneurysm development. Nature Communications 10:5046.

Pflanz, S., L. Hibbert, J. Mattson, R. Rosales, E. Vaisberg, J.F. Bazan, J.H. Phillips, T.K. McClanahan, R. de Waal Malefyt, and R.A. Kastelein. 2004. WSX-1 and Glycoprotein 130 Constitute a Signal-Transducing Receptor for IL-27. The Journal of Immunology 172:2225–2231.

Pflanz, S., J.C. Timans, J. Cheung, R. Rosales, H. Kanzler, J. Gilbert, L. Hibbert, T. Churakova, M. Travis, E. Vaisberg, W.M. Blumenschein, J.D. Mattson, J.L. Wagner, W. To, S. Zurawski, T.K. McClanahan, D.M. Gorman, J.F. Bazan, R. de Waal Malefyt, D. Rennick, and R.A. Kastelein. 2002. IL-27, a Heterodimeric Cytokine Composed of EBI3 and p28 Protein, Induces Proliferation of Naive CD4+ T Cells. Immunity 16:779–790.

Pietras, E.M., R. Lakshminarasimhan, J.M. Techner, S. Fong, J. Flach, M. Binnewies, and E. Passegué. 2014. Re-entry into quiescence protects hematopoietic stem cells from the killing effect of chronic exposure to type I interferons. J Exp Med 211:245–262.

Revuelta, M., and A. Matheu. 2017. Autophagy in stem cell aging. Aging Cell 16:912–915.

Reynaud, D., E. Pietras, K. Barry-Holson, A. Mir, M. Binnewies, M. Jeanne, O. Sala-Torra, J.P. Radich, and E. Passegué. 2011. IL-6 controls leukemic multipotent progenitor cell fate and contributes to chronic myelogenous leukemia development. Cancer Cell 20:661–673.

Ross, J.B., L.M. Myers, J.J. Noh, M.M. Collins, A.B. Carmody, R.J. Messer, E. Dhuey, K.J. Hasenkrug, and I.L. Weissman. 2024. Depleting myeloid-biased haematopoietic stem cells rejuvenates aged immunity. Nature 628:162–170.

Samusik, N., Z. Good, M.H. Spitzer, K.L. Davis, and G.P. Nolan. 2016. Automated mapping of phenotype space with single-cell data. Nat Methods 13:493–496.

Säwen, P., M. Eldeeb, E. Erlandsson, T.A. Kristiansen, C. Laterza, Z. Kokaia, G. Karlsson, J. Yuan, S. Soneji, P.K. Mandal, D.J. Rossi, and D. Bryder. 2018. Murine HSCs contribute actively to native hematopoiesis but with reduced differentiation capacity upon aging. eLife 7:e41258.

Schindelin, J., I. Arganda-Carreras, E. Frise, V. Kaynig, M. Longair, T. Pietzsch, S. Preibisch, C. Rueden, S. Saalfeld, B. Schmid, J.-Y. Tinevez, D.J. White, V. Hartenstein, K. Eliceiri, P. Tomancak, and A. Cardona. 2012. Fiji: an open-source platform for biological-image analysis. Nature Methods 9:676–682.

Seita, J., M. Asakawa, J. Ooehara, S. Takayanagi, Y. Morita, N. Watanabe, K. Fujita, M. Kudo, J. Mizuguchi, H. Ema, H. Nakauchi, and T. Yoshimoto. 2008. Interleukin-27 directly induces differentiation in hematopoietic stem cells. Blood 111:1903–1912.

Seyfried, A.N., J.M. Maloney, and K.C. MacNamara. 2020. Macrophages Orchestrate Hematopoietic Programs and Regulate HSC Function During Inflammatory Stress. Front Immunol 11:1499.

Siminovitch, L., J.E. Till, and E.A. McCulloch. 1964. Decline in colony-forming ability of marrow cells subjected to serial transplantation into irradiated mice. Journal of Cellular and Comparative Physiology 64:23–31.

Singh, S., B. Jakubison, and J.R. Keller. 2020. Protection of hematopoietic stem cells from stress-induced exhaustion and aging. Curr Opin Hematol 27:225–231.

Singh, S.K., S. Singh, S. Gadomski, L. Sun, A. Pfannenstein, V. Magidson, X. Chen, S. Kozlov, L. Tessarollo, K.D. Klarmann, and J.R. Keller. 2018. <em>Id1</em> Ablation Protects Hematopoietic Stem Cells from Stress-Induced Exhaustion and Aging. Cell Stem Cell 23:252–265.e258.

Stifter, S.A., N. Bhattacharyya, A.J. Sawyer, T.A. Cootes, J. Stambas, S.E. Doyle, L. Feigenbaum, W.E. Paul, W.J. Britton, A. Sher, and C.G. Feng. 2019. Visualizing the Selectivity and Dynamics of Interferon Signaling In Vivo. Cell Rep 29:3539–3550.e3534.

Sun, D., M. Zhang, and M. Shi. 2017. IL-27 limits neutrophil mediated pathology during pulmonary infection with Cryptococcus neoformans. The Journal of Immunology 198:123.128–123.128.

Villarino, A., L. Hibbert, L. Lieberman, E. Wilson, T. Mak, H. Yoshida, R.A. Kastelein, C. Saris, and C.A. Hunter. 2003. The IL-27R (WSX-1) is required to suppress T cell hyperactivity during infection. Immunity 19:645–655.

Weissman, I.L. 2000. Stem Cells: Units of Development, Units of Regeneration, and Units in Evolution. Cell 100:157–168.

Wirtz, S., I. Tubbe, P.R. Galle, H.J. Schild, M. Birkenbach, R.S. Blumberg, and M.F. Neurath 2006. Protection from lethal septic peritonitis by neutralizing the biological function of interleukin 27. Journal of Experimental Medicine 203:1875–1881.

Zhao, H.G., and M. Deininger. 2023. Always stressed but never exhausted: how stem cells in myeloid neoplasms avoid extinction in inflammatory conditions. Blood 141:2797–2812.

Zhu, J., J. Yu, A. Hu, J.Q. Liu, X. Pan, G. Xin, W.E. Carson, Z. Li, Y. Yang, and X.F. Bai. 2023. IL-27 Gene Therapy Induces Stat3-Mediated Expansion of CD11b+Gr1+ Myeloid Cells and Promotes Accumulation of M1 Macrophages in the Tumor Microenvironment. J Immunol 211:895–902.

